# A multi-enhancer hub at the *Ets1* locus controls T cell differentiation and allergic inflammation through 3D genome topology

**DOI:** 10.1101/2022.10.28.514213

**Authors:** Aditi Chandra, Sora Yoon, Michael F. Michieletto, Naomi Goldman, Emily K. Ferrari, Maria Fasolino, Leonel Joannas, Barbara L. Kee, Jorge Henao-Mejia, Golnaz Vahedi

## Abstract

Multi-enhancer hubs are spatial clusters of enhancers which have been recently characterized across numerous developmental programs. Yet, the functional relevance of these three-dimensional (3D) structures is poorly understood. Here we show that the multiplicity of enhancers interacting with the transcription factor *Ets1* is essential to control the precise expression level of this gene in response to cellular cues, and the failure to do so can lead to allergic diseases. Focusing on T cells as a model, we identified a highly connected multi-enhancer hub at the *Ets1* locus, comprising a noncoding regulatory element that is a hotspot for sequence variation associated with allergic diseases. We deleted this hotspot and found that the multi-enhancer connectivity is dispensable for T cell development but required for CD4^+^ T helper (Th1) differentiation in response to changes in the cytokine milieu. Mice lacking this hotspot are thus protected from Th1-mediated colitis but demonstrate an overt allergic response to house dust mites, a T cell-mediated response which is dampened by Th1 cells. Mechanistically, the multi-enhancer hub controls the expression level of *Ets1* that is dispensable for the active enhancer landscape but required for the Th1-specific genome topology through recruitment of CTCF. Together, we establish a paradigm for the functional and mechanistic relevance of multi-enhancer hubs controlling cellular competence to respond specifically to an inductive cue.

## Introduction

The spatiotemporal control of gene-expression programs relies on nuclear organization of noncoding regulatory elements called enhancers. Recent advances in genomics and imaging technologies attest to the formation of spatial clusters of enhancers, called interchangeably as multi-enhancer hubs^1–8^, 3D cliques^9,10^, *cis*-regulatory domains^11^, interacting triplets^12^, connected gene communities^13^, or architectural stripes^14,15^. Despite numerous efforts profiling multi-enhancer interactions across diverse developmental programs, the functional relevance of these structures is poorly understood.

We reasoned that searching for disease-associated sequence variations that overlap regulatory elements converging into multi-enhancer hubs can put forward candidates for functional and mechanistic investigations of multi-enhancer hubs. We focused on T cells as a model and mathematically defined the higher-order structure of multi-enhancer interactions in mouse T cells^10^. Our unbiased strategy revealed that the *Ets1-Fli1* locus is among the most hyperconnected regions, forming a multi-enhancer hub that is also conserved in human T cells. Here we found that while numerous single-nucleotide-polymorphisms (SNPs) associated with immune-mediated diseases were distributed across the human *ETS1*-*FLI1* locus, a noncoding element within this multi-enhancer hub was found to be a hotspot for SNPs associated with type 2 immune diseases including allergy, asthma, and atopic dermatitis^16,17^. Although complete ablation of *Ets1* is linked to immune-mediated diseases^18,19^, mechanisms through which noncoding sequence variation at this locus contributes to allergic responses are unknown.

ETS1 and FLI1 are ETS family transcription factors containing a conserved winged helix-turn-helix DNA-binding domain^20,21^. The members of this transcription factor family of proteins bind to a consensus DNA sequence containing a core GGA(A/T) motif^22^. ETS1, which has diverse roles in multiple biological processes^23^, is most highly expressed in lymphoid organs^24,25^ and is required for the development of T cells, natural killer (NK) cells, and innate lymphoid cells^18,19,23,24,26–29^. FLI1 is not as highly expressed as ETS1 in lymphoid organs, yet its contribution to effector CD8^+^ T cell function has been described^30^.

We performed *in vivo* deletion of the regulatory node at the *Ets1* locus in mice using the CRISPR–Cas9 system. We found that the genetic deletion of this regulatory element does not affect T cell development but impairs CD4^+^ T helper 1 (Th1) cell differentiation, leading to protection against colitis, a Th1-mediated disease, and an overt allergic response, a type 2 immune response dampened by Th1 cells. Our molecular, optical, and cellular analyses propose a mechanism through which the multiplicity of enhancer interactions at the *Ets1* locus controls the distinct expression level of *Ets1* in response to changes in the cytokine environment which in turn is required for the recruitment of CTCF to specify three-dimensional (3D) nuclear organization of Th1 cells. Considering that Th1 differentiation is a critical mechanism by which type 2 immune responses are dampened^31–33^, our findings imply the molecular processes through which sequence variation within noncoding elements at the *Ets1* locus predisposes individuals to allergic responses. These findings establish a paradigm for understanding the importance of multi-enhancer hubs in dynamically regulating gene expression in response to changes in cellular environment.

## Results

### Multi-enhancer connectivity at T cell-lineage genes

We aimed to identify multi-enhancer hubs in T cells in an unbiased manner and leveraged human genetics to ascertain whether sequence variation within the top densely connected multi-enhancer hubs in T cells is linked to immune-mediated diseases. We reasoned that mapping multi-enhancer interactions in thymocytes, which are T cells before any antigen exposure can delineate critical regulatory units shared across T cell populations. Hence, we mapped enhancer interactions in double-positive (DP) thymocytes and used H3K27ac HiChIP^10^, which is a protein-centric assay for 3D mapping of enhancer interactions^34,35^. We algorithmically searched for groups of densely connected multi-enhancers, which have also been referred to as 3D cliques^9,10^ (Fig. 1a). The degree of enhancer connectivity was asymmetrical, reminiscent of asymmetrical histone acetylation at super-enhancers^36,37^. Although most multi-enhancer hubs contained less than 7 interacting elements, 2,372 regulatory elements spatially converged into 115 “hyperconnected” multi-enhancer hubs (Table S1). Super-enhancers were significantly enriched in hyperconnected hubs, suggesting extensive spatial connectivity among highly acetylated genomic elements (Fig. 1b). Although not statistically significant, around 20% of hyperconnected hubs overlapped with annotated noncoding RNAs (Fig. S1a) and approximately 70% of hyperconnected hubs were characterized as architectural stripes^14,15^, which are thought to form through the loop extrusion process when a loop anchor interacts with the entire domain at a high frequency (Fig. S1b). Overall, genes associated with T cell biology, including “T cell receptor signaling pathway” and “adaptive immune system”, were highly enriched at hyperconnected hubs (Fig. 1c), thus suggesting that our analytical approach can prioritize regions harboring genes critical for T cells.

**Figure 1:**
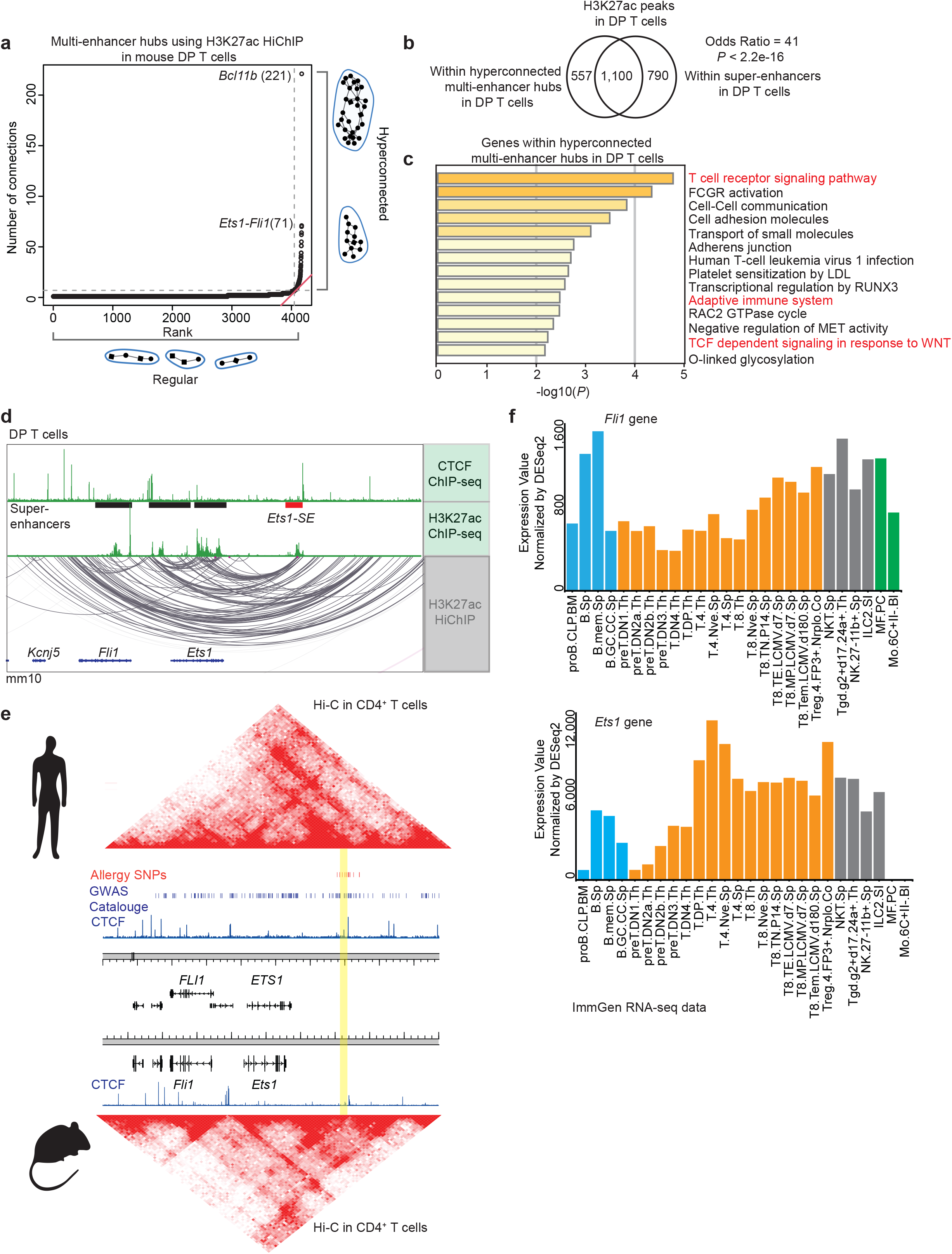
Exceptional enhancer connectivity at the *Ets1-Fli1* locus. **a,** Plot depicts ranking versus number of connections in multi-enhancer hubs also referred to as 3D cliques^9,10^ detected in double-positive (DP) T cells using H3K27ac HiChIP measurements generated in our previous study^10^. H3K27ac HiChIP measurements in two biological replicates were generated, and only reproducible interactions were used for this analysis. Multi-enhancer hubs are plotted in an ascending order of their total connectivity. Hyperconnected multi-enhancer hubs are defined as the ones above the elbow of the total connectivity ranking. Top two hyperconnected multi-enhancer hubs *Bcl11b* and *Ets1-Fli1* are labeled. Number of interactions in each hub is provided in parenthesis. **b**, Venn-diagram depicts the overlaps of genomic regions within hyperconnected multi-enhancer hubs detected using H3K27ac HiChIP when compared to super-enhancers defined by H3K27ac ChIP-seq in DP T cells. Super-enhancers were defined as described before^36^ based on H3K27ac ChIP-seq performed in three biological replicates in DP T cells. Odds ratio and Fisher’s exact test were used for statistical analysis. **c,** Bar-plot demonstrates the significance of gene-ontology terms enriched in genes encompassing hyperconnected 3D cliques. Metascape^58^ was used for gene-ontology analysis. Terms relating to immune response pathways are marked in red. **d,** The genome browser view demonstrates H3K27ac and CTCF ChIP-seq, as well as H3K27ac HiChIP 3D interactions at the *Ets1-Fli1* locus in DP T cells. H3K27ac HiChIP measurements in two biological replicates were generated and only reproducible interactions were used for visualization. The 25 kbp super-enhancer which is the focus of this study is called *Ets1*-SE and is marked in red. **e,** Heatmaps demonstrate contact-frequency maps measured by ultra-high resolution Hi-C in CD4^+^ T cells in humans and mice. ChIP-seq tracks demonstrate CTCF binding in CD4^+^ T cells in humans and DP T cells in mice. SNPs associated with asthma and allergic diseases highlighted in red were curated from the GWAS catalogue and meta-analysis studies for allergy, asthma, and atopic dermatitis^16,17^. Blue bars demonstrate statistically significant GWAS SNPs associated with diverse traits. Homologous locus to the *Ets1-*SE is highlighted in yellow. **f,** Bar plots demonstrate *Ets1* and *Fli1* gene expression levels in B cells (blue), T cells (orange), NK cells (gray) and macrophages (green) using ImmGen RNA-seq normalized by DESeq2.

The top hyperconnected locus in thymocytes encompassed the transcription factor *Bcl11b* (Fig. 1a). Bcl11b is expressed in T cell progenitors and controls a T-lineage-specific program by suppressing alternative cell fates^38^. One of the major enhancer elements in this hyperconnected locus repositions from the peripheral lamina to the nuclear interior, a process which is required for T cell development and lymphomas^39^. The identification of *Bcl11b* as the most hyperconnected locus in T cells highlights the sensitivity of our analytical approach in identifying genes with key biological roles in this lineage. Yet, it is unknown whether hyperconnectivity of enhancers at other genomic loci is essential for T cell functional competence.

### Exceptional enhancer connectivity at a type 2 immune disease-associated risk locus

The second most hyperconnected multi-enhancer hub was detected at the locus encompassing two ETS-family transcription factors, *Ets1* and *Fli1*, and demonstrated conserved chromatin folding patterns between human and mouse T cells (Fig. 1a,d,e). Whereas numerous SNPs associated with diverse diseases were distributed within the ∼700kbp region encompassing *ETS1* and *FLI1* in the human genome, 12 independent variants associated with type 2 immune diseases, namely self-reported allergy^16^, asthma^16^, and atopic dermatitis^17^, concentrated within a 25 kbp DNA segment that is ∼250 kbp downstream of the *ETS1* promoter (SNPs in red, segment marked in yellow, Fig. 1e). This regulatory segment was a major node of enhancer connectivity, overlapped with an annotated noncoding RNA, and scored as a super-enhancer, and thus is hereafter referred to as “*Ets1-*SE*”* (red block, Fig. 1d).

The unusual clustering of type 2 immune disease associated SNPs around the *Ets1-*SE element provided the rationale for testing the functional relevance of this regulatory sequence. Hence, we generated a novel mouse strain in which the 25 kbp *Ets1-*SE is deleted on the C57BL/6J background (red block, Fig. 1d). In contrast with *Fli1*, *Ets1* expression increases during T cell development (Fig. 1f). Together, the high expression of *Ets1* during development along with the hyperconnectivity of the *Ets1* locus in T cells was the motivation to study the effect of the *Ets1-*SE deletion on T cell development and function.

### Multi-enhancer interactions in the *Ets1* locus control CD4^+^ T helper 1 differentiation

Since ETS1 is required for the T cell lineage^23^, we assessed the effect of the *Ets1-*SE deletion on T cell development. Surprisingly, T cell development remained intact in *Ets1*-SE^−/−^ mice as the proportion of cells in the different thymic T cell developmental stages as well as the numbers of natural regulatory T cells (Tregs) were not significantly different when compared to wildtype mice (Figs. 2a and S2a-b). At steady-state, the majority of CD4^+^ and CD8^+^ T cell populations in peripheral tissues remained intact in *Ets1*-SE^−/−^ mice when compared to wildtype animals (Fig. S2c-e), apart from a moderate reduction in CD4^+^ T cell numbers in the lungs (Fig. 2b). Together, unlike the *Bcl11b* locus^39^, perturbation of the hyperconnected *Ets1* locus was tolerated during T cell development.

**Figure 2:**
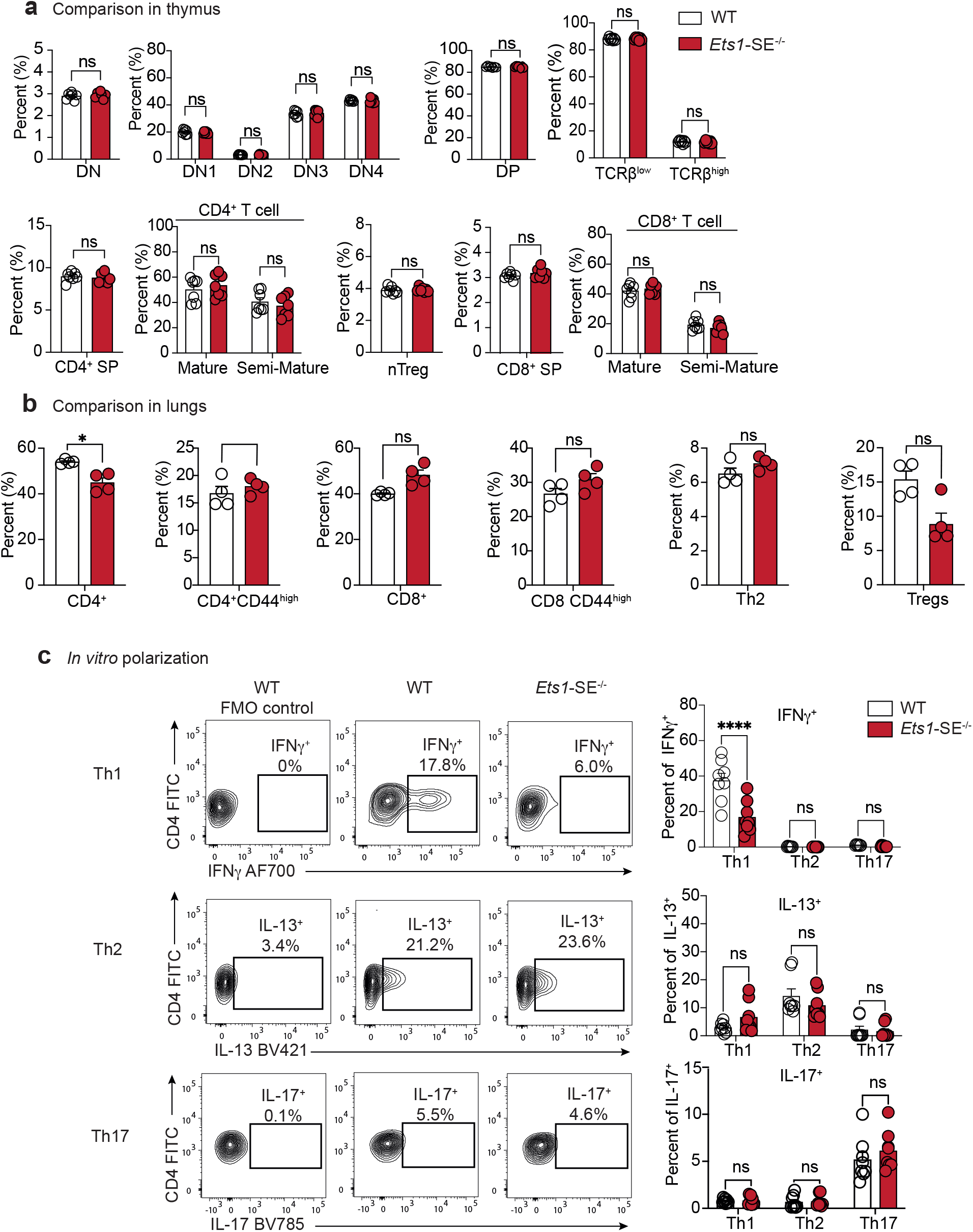
*Ets1*-SE is dispensable for thymic T cell generation but is required for CD4^+^ Th1 differentiation. **a,** Plots demonstrate percentage of cells defined by flow cytometry analysis in the thymus. Quantification in the thymus is defined as: Double Negative (DN; TCRβ^+^, CD4^−^, CD8^−^), DN1 (TCRβ^+^, CD4^−^, CD8^−^, CD44^+^, CD25^−^), DN2 (TCRβ^+^, CD4^−^, CD8^−^, CD44^+^, CD25^+^), DN3 (TCRβ^+^, CD4^−^, CD8^−^, CD44^−^, CD25^+^), DN4 (TCRβ^+^, CD4^−^, CD8^−^, CD44^−^, CD25^−^), TCRβ^low^ (Live, TCRβ^low^), TCRβ^high^ (Live, TCRβ^high^), Double Positive (DP; TCRβ^+^, CD4^+^, CD8^+^), single-positive CD4^+^ T cells (CD4SP; TCRβ^+^, CD4^+^, CD8^−^), semi mature CD4^+^ T cells (TCRβ^+^, CD4^+^, CD24^+^), Mature CD4^+^ T cells (TCRβ^+^, CD4^+^, CD24^−^), single positive CD8^+^ T cells (CD8SP; TCRβ^+^, CD4^−^, CD8^+^), semi-mature CD8^+^ T cells (TCRβ^+^, CD8^+^, CD24^+^), mature CD8^+^ T cells (TCRβ^+^, CD8^+^, CD24^−^), immature single positive CD8^+^ T cells (iSP CD8; TCRβ^low^, CD8^+^, CD24^high^) and thymic regulatory CD4^+^ T cells (TCRβ^+^, CD4^+^, FoxP3^+^) from age matched wildtype and *Ets1-*SE^−/−^ female mice. All cell populations were pre-gated on SSC-A/FSC-A, Singlets and Viability^−^ (Live) cells and complete gating strategy used to identify each population is displayed in **Figure S2a**. Data are representative of three independent experiments. Each dot represents an individual mouse (wildtype; n=7 - *Ets1-*SE^−/−^; n=7). Error bars = SEM; and *P*: ns = not significant, (DN, DP, nTregs, CD4^+^/CD8^+^ SP: Mann-Whitney U test; DN1-DN4, TCRβ^low/high^, Semi-Mature/Mature CD4^+^/CD8^+^: Two-way ANOVA with multiple comparisons and Bonferroni correction). **b,** Plots demonstrate percentages of cells defined by flow cytometry analysis in the lung Frequencies at steady state of lungs parenchyma CD4^+^ T cells (TCRβ^+^, CD4^+^), activated CD4^+^ T cells (TCRβ^+^, CD4^+^, CD44^+^), CD8^+^ T cells (TCRβ^+^, CD8^+^), activated CD8^+^ T cells (TCRβ^+^, CD8^+^, CD44^+^), CD4^+^ Th2 cells (TCRβ^+^, CD4^+^, GATA-3^+^) and regulatory CD4^+^ T cells (TCRβ^+^, CD4^+^, Foxp3^+^) from age-matched wildtype and *Ets1-*SE^−/−^ male mice. All populations were pre-gated on SSC-A/FSC-A, Singlets and Viability^−^ (Live) cells and complete gating strategy used to identify each population is displayed in **Figure S2b**. Data are representative of two independent experiments. Each dot represents an individual mouse (wildtype; n=4 - *Ets1-SE*^−/−^; n=4). Error bars = SEM; and *P*: ns = not significant, * =*P*≤0.05 (Mann-Whitney U test). **c,** (left) Representative flow cytometry contour plot of naive CD4^+^ T cells from wildtype or *Ets1-* SE^−/−^ mice cultured under Th1, Th2 or Th17 polarizing conditions for 6 days. Unstained wildtype cells are shown for each polarizing condition as a negative control (wildtype FMO Control). (right) Frequencies of Th1 (IFNγ^+^), Th2 (IL-13^+^) or Th17 (IL-17^+^) cytokines producing CD4^+^ T cells cultured under Th1, Th2 or Th17 polarizing conditions for 6 days. All populations were pre-gated on SSC-A/FSC-A, Singlets and Viability^−^ (Live), TCRβ^+^ and CD4^+^ cells. Two independent experiments were pooled and were repeated five times. Each dot represents an individual mouse (wildtype; n=8 - *Ets1-SE*^−/−^; n=8). Error bars = SEM; and *P*: ns = not significant, * = *P* ≤0.05, ** = *P* ≤0.01, *** = *P* ≤0.0005, **** = *P* ≤0.0001 (Two-way ANOVA with multiple comparisons and Bonferroni correction).

The concentration of genetic variants associated with dysregulated type 2 immune responses within *Ets1*-SE implies a link between this region and CD4^+^ T differentiation, which is a process mediated by changes in the extracellular cytokine milieu evoked by pathogenic stimuli^40^. Therefore, we next aimed to explore whether perturbation in the *Ets1* multi-enhancer hub affected the capacity of CD4^+^ T cells to differentiate into distinct T helper functional subsets namely Th1, Th2, and Th17 populations when exposed to their respective polarizing milieu *in vitro*^31^. Strikingly, naïve CD4^+^ T cells from *Ets1-*SE^−/−^ mice had a significantly reduced capacity to differentiate into interferon-gamma (IFNγ) producing Th1 cells when compared to wildtype CD4^+^ T cells (Fig. 2c). In contrast, Th2 and Th17 differentiation measured by IL-13 and IL-17 expression, respectively, remained comparable between wildtype and *Ets1-*SE^−/−^ CD4^+^ T cells (Fig. 2c). Thus, the *Ets1-*SE element controls CD4^+^ Th1 differentiation *in vitro*.

### *Ets1*-SE element promotes the development of severe colitis *in vivo*

To determine if *Ets1*-SE deletion is associated with impaired Th1 differentiation *in vivo*, we tested the capacity of *Ets1*-SE^−/−^ CD4^+^ T cells to induce colitis using a well-established adoptive transfer Th1-driven colitis model^41^. To do so, we adoptively transferred CD4^+^CD45RB^high^ T cells from wildtype or *Ets1*-SE^−/−^ mice into *Rag1*^−/−^ mice, which lack mature B and T lymphocytes, and monitored their body weights as well as colitis development for 6 weeks (Fig. 3a). As expected^41^, *Rag1*^−/−^ mice that received wildtype CD4^+^ T cells lost ∼10% of their initial body weights by week 6 post-T cell transfer while *Rag1*^−/−^ mice that received *Ets1*-SE^−/−^ CD4^+^ T cells steadily maintained their body weights at significantly higher levels when compared to *Rag1*^−/−^ mice that received wildtype cells (Fig. 3b). In concordance, *Rag1*^−/−^ mice that received *Ets1*-SE^−/−^ CD4^+^ T cells had significantly longer colon lengths and reduced severity in colonic histopathology when compared to *Rag1*^−/−^ recipient of wildtype T cells, thus indicating that Th1-driven colon inflammation was drastically reduced in presence of *Ets1-*SE*^−/−^* CD4^+^ T cells (Figs. 3c-e and S3a).

**Figure 3:**
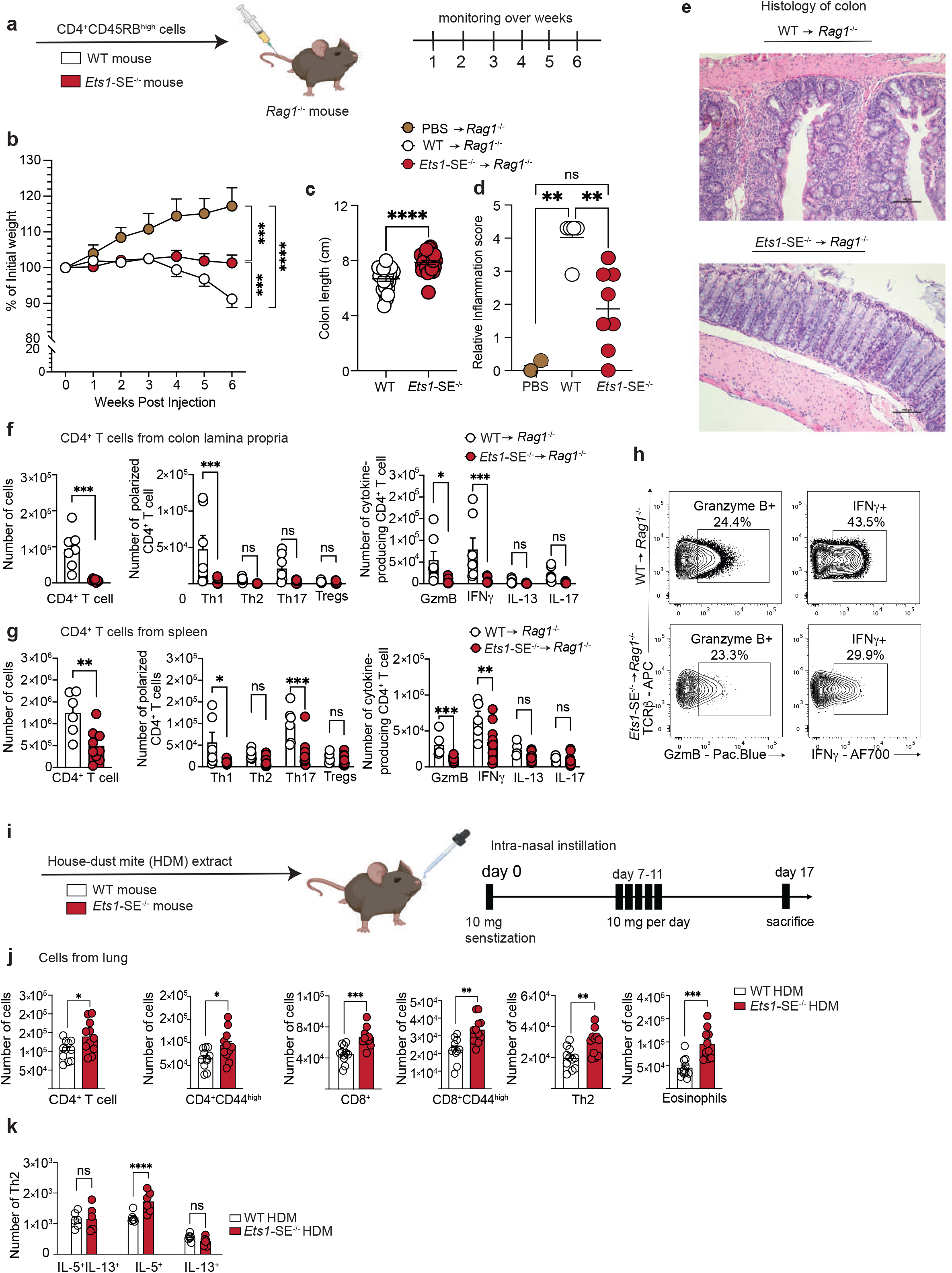
*Ets1*-SE deletion limits Th1-mediated inflammation *in vivo*. **a,** Schematic representation of the CD45RB^High^-induced colitis model. 1×10^6^ FACS sorted TCRβ^+^, CD4^+^, CD45RB^High^ naïve CD4^+^ T cells from wildtype or *Ets1-*SE^−/−^ were transferred into *Rag1*^−/−^ recipients to induce colitis. Weight was monitored once a week for six weeks post injection. **b,** Weight loss tracking of *Rag1*^−/−^ mice that received either 1×10^6^ FACS sorted TCRβ^+^, CD4^+^, CD45RB^High^ naïve wildtype or *Ets1-*SE^−/−^ CD4^+^ T cells as compared to PBS-injected animals (controls) during 6 weeks post transfer. Weight was measured once per week. Three independent experiments were pooled and repeated four times. Dots represent the mean of individual mouse (PBS -> *Rag1*^−/−^; n= 5; wildtype -> *Rag1*^−/−^; n=25 and *Ets1-SE*^−/−^ -> *Rag1*^−/−^; n=27). Error bars = SEM; and *P*: ns = not significant, * = *P*≤0.05, ** = *P*≤0.01, *** = *P*≤0.0005, **** = *P*≤0.0001 (Two-way ANOVA; Mixed-effect REML model with Fisher’s LSD Test). **c,** Quantification of colon length (cm) of *Rag1*^−/−^ mice that received either 1×10^6^ FACS sorted TCRβ^+^, CD4^+^, CD45RB^High^ naïve wildtype or *Ets1-*SE^−/−^ CD4^+^ T cells at week 6 post transfer. Three independent experiments were pooled and repeated four times. Dots represent the mean of individual mouse (wildtype -> *Rag1*^−/−^; n=25 and *Ets1-SE*^−/−^ -> Rag1^−/−^; n=27). Error bars = SEM; and *P*: ns = not significant, * = *P* ≤0.05, ** = *P* ≤0.01, *** = *P* ≤0.0005, **** = *P* ≤0.0001 (Mann-Whitney U Test). **d,e** Quantification of histological score of paraffin-embedded colon rolls from *Rag1*^−/−^ mice that received 1×10^6^ FACS sorted TCRβ^+^, CD4^+^, CD45RB^High^ naïve CD4^+^ T cells from either wildtype or *Ets1-*SE^−/−^ mice at week 6 post transfer and as compared to PBS-injected mice (control). Histological sections were obtained from two independent experiments and were scored in a blinded manner. Each dot represents an individual mouse (PBS -> *Rag1*^−/−^; n= 2, wildtype -> *Rag1*^−/−^; n=5 and *Ets1-SE*^−/−^ -> *Rag1*^−/−^; n=8). Error bars = SEM; and *P*: ns = not significant, * = *P*≤0.05, ** = *P* ≤0.01, *** = *P* ≤0.0005, **** = *P* ≤0.0001 (One-way ANOVA with multiple comparisons and Bonferroni correction). **e**, Histological section of colon rolls stained with H&E of *Rag1*^−/−^ mice that received either 1×10^6^ FACS sorted TCR^+^, CD4^+^, CD45RB^High^ naïve wildtype or *Ets1-*SE^−/−^ CD4^+^ T cells 6 weeks post transfer. Scale = 100μm; Magnification = 100X. **f,** (left) Quantification of infiltrating colon lamina propria (cLP) colitogenic CD4^+^ T cells of *Rag1*^−/−^ mice that received either 1×10^6^ FACS sorted TCRβ^+^, CD4^+^, CD45RB^High^ naïve wildtype or *Ets1-* SE^−/−^ CD4^+^ T cells at week 6 post transfer. (middle) Quantification of Th1 (T-bet^+^), Th2 (GATA-3^+^), Th17 (RORγ-t^+^) and Tregs (FoxP3^+^) CD4^+^ T cells among infiltrating cLP CD4^+^ T cells of *Rag1*^−/−^ mice that received either 1×10^6^ FACS sorted TCRβ^+^, CD4^+^, CD45RB^High^ naïve wildtype or *Ets1-* SE^−/−^ CD4^+^ T cells at week 6 post transfer. (right) Quantification of Th1 (IFNγ^+^), Th2 (IL-13^+^), Th17 (IL-17^+^) and Granzyme B (GzmB^+^) producing CD4^+^ T cells among infiltrating cLP CD4^+^ T cells of *Rag1*^−/−^ mice that received either 1×10^6^ FACS sorted TCRβ^+^, CD4^+^, CD45RB^High^ naïve wildtype or *Ets1-*SE^−/−^ CD4^+^ T cells at week 6 post transfer. Two independent experiments were pooled. Dots represent an individual mouse (*Ets1-SE*^+/+^ -> *Rag1*^−/−^; n=7 and *Ets1-SE*^−/−^ -> *Rag1*^−/−^; n=7). Error bars = SEM; and *P*: ns = not significant, * = *P* ≤0.05, ** = *P* ≤0.01, *** = *P* ≤0.0005, **** = *P*≤0.0001 (CD4^+^ T cells: Mann-Whitney U Test; Th1/Th2/Th17/Treg and cytokines production: Two-way ANOVA with multiple comparisons and Bonferroni correction). **g,** (left) Quantification CD4^+^ T cells in the spleen of *Rag1*^−/−^ mice that received either 1×10^6^ FACS sorted TCRβ^+^, CD4^+^, CD45RB^High^ naïve wildtype or *Ets1-*SE^−/−^ CD4^+^ T cells at week 6 post transfer. (middle) Quantification in the spleen of Th1 (T-bet^+^), Th2 (GATA-3^+^), Th17 (RORγ-t^+^) and Tregs (FoxP3^+^) CD4^+^ T cells of *Rag1*^−/−^ mice that received either 1×10^6^ FACS sorted TCRβ^+^, CD4^+^, CD45RB^High^ naïve wildtype or *Ets1-*SE^−/−^ CD4^+^ T cells at week 6 post transfer. (right) Quantification of *ex vivo* stimulated splenic Th1 (IFNγ^+^), Th2 (IL-13^+^), Th17 (IL-17^+^) and Granzyme B (Gzm B^+^) producing CD4^+^ T cells of *Rag1*^−/−^ mice that received either 1×10^6^ FACS sorted TCRβ^+^, CD4^+^, CD45RB^High^ naïve wildtype or *Ets1-*SE^−/−^ CD4^+^ T cells at week 6 post transfer. Data are a representative of one experiment which was repeated three times. Dots represent an individual mouse (wildtype -> *Rag1*^−/−^; n=7 and *Ets1-*SE^−/−^ -> *Rag1*^−/−^; n=7). Error bars = SEM; and p-values: ns = not significant, * = *P*≤0.05, ** = *P* ≤0.01, *** = *P* ≤0.0005, **** = *P* ≤0.0001 (CD4^+^ T cells: Mann-Whitney U Test; Th1/Th2/Th17/Treg and cytokines production: Two-way ANOVA with multiple comparisons and Bonferroni correction). **h,** Representative flow cytometry contour plot of Granzyme B- and IFNγ-producing *ex vivo* stimulated cLP CD4^+^ T cells of *Rag1*^−/−^ mice that received either 1×10^6^ FACS sorted TCRβ^+^, CD4^+^, CD45RB^High^ naïve wildtype or *Ets1-*SE^−/−^ CD4^+^ T cells at week 6 post transfer. **i,** Schematic representation of the house dust mite (HDM) extract challenge. Arrows represent days by which intranasal HDM was administered. Mice were euthanized 16 days after the initial sensitization and immune cell infiltration was checked. Two independent experiments were used to measure immune cell infiltration in the lung parenchyma. **j,** Quantification of lung parenchyma infiltrating CD4^+^ T cells (TCRβ^+^, CD4^+^), activated CD4^+^ T cells (TCRβ^+^, CD4^+^, CD44^+^), CD8^+^ T cells (TCRβ^+^, CD8^+^), activated CD4^+^ T cells (TCRβ^+^, CD8^+^, CD44^+^), CD4^+^ Th2 cells (TCRβ^+^, CD4^+^, GATA-3^+^) and eosinophils (MHC-II^−^, Siglec-F^+^) from wildtype or *Ets1-*SE^−/−^ mice 16 days after HDM challenge. All cells were pre-gated on SSC-A/FSC-A, Singlets, Live cell (Viability^−^). Two independent experiments were pooled. Each Dot represents an individual mouse (wildtype n=11 and *Ets1-SE*^−/−^ n=11). Error bars = SEM; and ns = not significant, * = *P* ≤0.05, ** = *P* ≤0.01, *** = *P* ≤0.0005, **** = *P* ≤0.0001 (Mann-Whitney). **k,** Quantification of Th2 cells numbers from wildtype or *Ets1-*SE^−/−^ mice producing IL-5, IL-13 or IL-5/IL-13 four hours after PMA / Ionomycin stimulation. All cells were pre-gated on SSC-A/FSC-A, Singlets, Live cell (Viability^−^), CD3^+^, CD90.2^+^, CD4^+^, ST2^+^. Data are representative of one independent experiment and repeated twice. Each dot represents an individual mouse (wildtype n=6 and *Ets1-*SE^−/−^ n=6). Error bars = SEM; and *P*: ns = not significant, * = *P* ≤0.05, ** = *P* ≤0.01, *** = *P* ≤0.0005, **** = *P* ≤0.0001 (Two-way ANOVA with multiple comparison and Bonferroni correction).

To evaluate the functional profiles of CD4^+^ T cells in *Rag1*^−/−^ recipient mice, we isolated CD4^+^ T cells from the colon and spleen of these animals at least 6 weeks after the CD4^+^ CD45RB^high^ T cell transfers and measured the frequency and numbers of Th1, Th2, Th17, and Treg populations. In concordance with a decrease in capacity of *Ets1*-SE^−/−^ CD4^+^ T cells to differentiate into IFNγ-producing Th1 cells, *Rag1*^−/−^ mice that received wildtype CD4^+^ T cells had significantly higher total numbers of CD4^+^ T cells in the colon lamina propria and spleen as compared to *Rag1*^−/−^ mice that received *Ets1*-SE^−/−^ CD4^+^ T cells (Fig. 3f,g). Moreover, numbers of Granzyme B- and IFNγ-producing CD4^+^ T cells were lower in *Ets1-SE^−/−^* injected *Rag1^−/−^* animals as compared to wildtype (Fig. 3h), while numbers of Th2 or Tregs remained comparable (Fig. 3f-h). Hence, *Ets1-*SE in CD4^+^ T cells is specifically required for Th1 differentiation *in vivo* in the context of a Th1-induced colitis model.

### *Ets1*-SE deletion induced type 2 immune responses *in vivo*

IFNγ production by Th1 cells is a critical mechanism that dampens Th2 responses^31–33^. Since several SNPs associated with allergic responses are clustered around the *Ets1*-SE element, we postulated that compromised Th1 differentiation in *Ets1*-SE^−/−^ mice can lead to enhanced allergic responses *in vivo*. Hence, we challenged wildtype and *Ets1*-SE^−/−^ mice with House Dust Mite (HDM) extracts for 5 consecutive days after an initial exposure and quantified immune cell infiltration and type 2 cytokine production in the lungs 17 days after the initial exposure (Fig. 3i). We found a significantly higher number of infiltrating CD4^+^ and CD8^+^ T cells in the lungs of *Ets1*-SE^−/−^ animals compared to wildtype counterparts (Fig. 3j). Notably, we observed dramatically increased numbers of eosinophils and Th2 cells in HDM-challenged *Ets1*-SE^−/−^ mice, a key signature of over-active type 2 responses (Figs. 3j and S3b). Indeed, *Ets1*-SE^−/−^ mice also showed a significantly higher number of IL-5^+^ producing Th2 cells during HDM-induced allergic airway inflammatory responses (Fig. 3k). Thus, the *Ets1-*SE region is required to dampen Th2 responses by promoting optimal differentiation of Th1 cells, providing a mechanistic link between the human genetic association of this locus with asthma and allergic diseases.

### Reorganization in the multi-enhancer hub after the *Ets1*-SE deletion

Thus far, we showed that the *Ets1*-SE element is required for optimal Th1 differentiation *in vitro* and *in vivo* and the genetic deletion of this element leads to allergic responses (Figs. 2-3). Yet, the molecular processes through which the *Ets1*-SE element controls CD4^+^ T cell differentiation remain unknown. We first assessed how the multi-enhancer hub was affected in the absence of the *Ets1*-SE element at a single-cell resolution. Using the flexible, scalable, and high-resolution ‘‘Oligopaint’’ DNA fluorescence *in situ* hybridization (FISH) approach^42,43^, we painted 3 anchors of the *Ets1* multi-enhancer hub by designing probes spanning ∼50kbp regions. We chose to probe multi-enhancer interactions including an enhancer upstream of *Ets1* (E1), the *Ets1* promoter and gene-body (E2), and an enhancer downstream of the *Ets1*-SE element (E3), where all elements were intact in both wildtype and *Ets1*-SE^−/−^ genomes (Fig. 4a). To evaluate the early effect of T cell activation, we performed these Oligopaint experiments in activated CD4^+^ T cells, only 18 hours after T cell receptor activation. On average, the spatial distance between probe-pairs reduced in *Ets1*-SE^−/−^ compared with wildtype CD4^+^ T cells (Figs. 4b and S4a-b). Strikingly, studying the individual alleles revealed that in the absence of *Ets1*-SE, the frequency of individual cells whose upstream enhancer E1 was spatially proximal (<0.5 μm) to the *Ets1* promoter E2 increased (Figs. 4c and S4c). Moreover, multi-enhancer interactions (or 3D cliques) where all three regulatory elements converged into a hub in the same cell were detected in 2x more alleles in *Ets1*-SE^−/−^ compared with wildtype T cells, suggesting that the *Ets1*-SE deletion can rewire the *Ets1* locus (Fig. 4d). Representative cells from wildtype and *Ets1*-SE^−/−^ mice further visualized the formation of multi-enhancer interactions (Fig. 4e). Together, the *Ets1*-SE deletion led to reorganization of the multi-enhancer hub, suggesting the pliability of the genome at these highly interacting loci and a potential compensatory mechanism in the absence of *Ets1*-SE.

**Figure 4.**
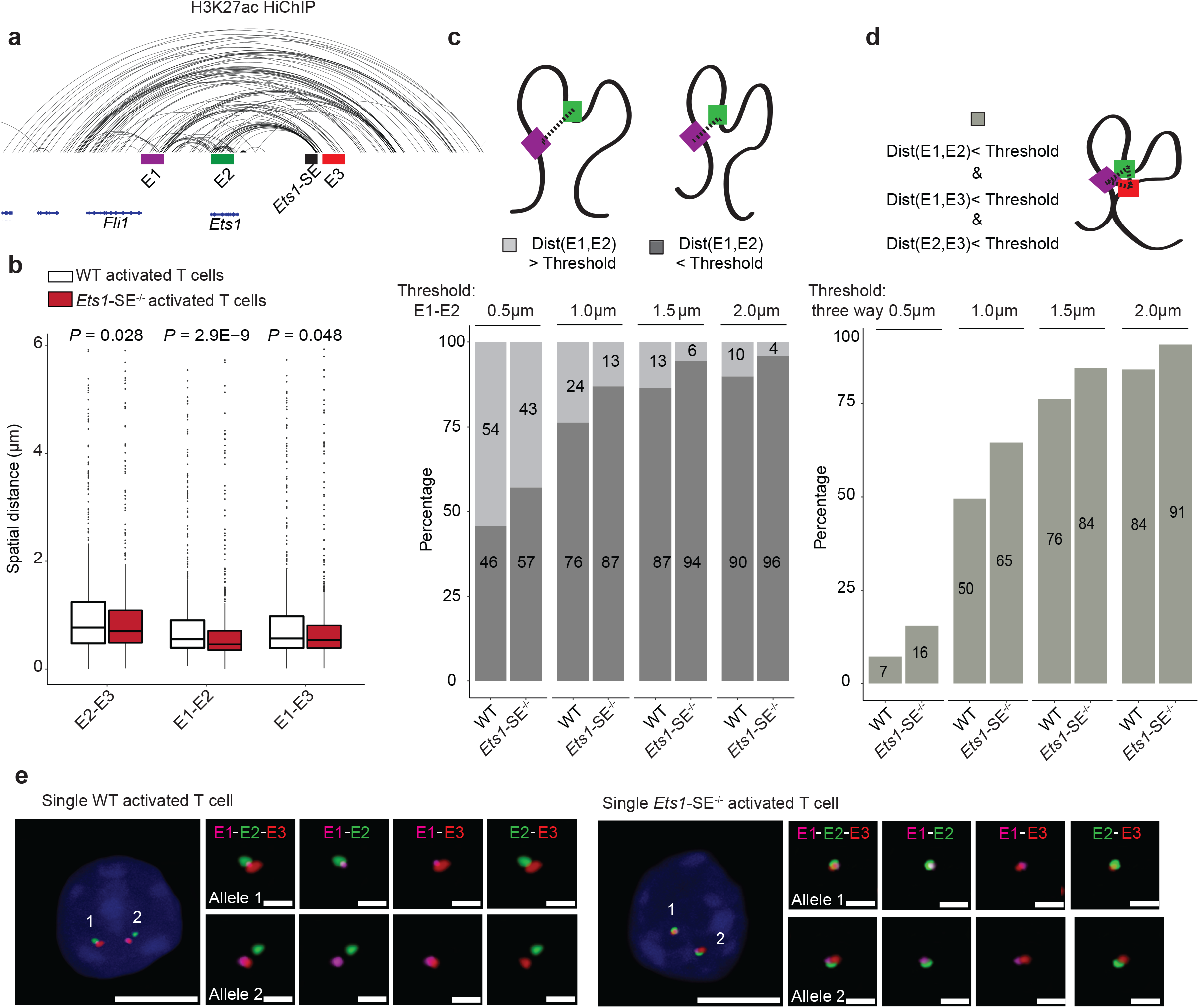
Reorganization of the multi-enhancer hub in the absence of the *Ets1*-SE element. **a**, H3K27ac-HiChIP contacts in DP T cells at the *Ets1* locus, with E1 (enhancer upstream of *Ets1*, magenta), E2 (*Ets1* promoter and gene-body, green), and E3 (enhancer downstream of *Ets1*-SE, red) representing the three independent 50 kbp genomic regions for which Oligopaint probes were designed. The *Ets1*-SE element deleted in mice is depicted in black. **b**, Box plots of the spatial distance formed between E1 and E2 probes (KS: Kolmogorov-Smirnov test & MW: Mann-Whitney test). Oligopaint 3D FISH^42,59^ in more than 500 activated CD4^+^ T cells from the spleen were imaged using widefield microscopy from one wildtype and one *Ets1*-SE^−/−^ mouse. CD4^+^ T cells 18 hours after activation with T cell receptor signaling per genotype were used. Spots for each probe in Oligopaint 3D FISH data were analyzed in a semi-manual manner (Methods). **c**, Bar plots demonstrating percentage of activated CD4^+^ T cells where spatial distance between genomic regions corresponding to E1 and E2 probes is less than a selected threshold (0.5μm, 1μm, 1.5μm, or 2μm) (dark gray) compared with cells whose E1 to E2 distance is greater than a selected threshold (light gray). **d**, Bar plots demonstrating percentage of activated CD4^+^ T cells forming a multi-enhancer hub or a 3D clique where three-way spatial distances between E1-E2, E1-E3, and E2-E3 probes are all less than a selected threshold (0.5μm, 1μm, 1.5μm, or 2μm). **e**, Representative images of the Oligopaint FISH probes in one wildtype and one *Ets1*-SE^−/−^ activated CD4^+^ T cell, with magnification of each allele. DAPI: blue, E1: magenta, E2: green, E3: red. Scale-bar in whole cell: 5μm and scale-bar in magnification of allele: 1μm.

### Transcriptional outputs of Th1 cells depend on *Ets1*-SE

Considering the strong *in vivo* phenotype and genome reorganization in the absence of *Ets1*-SE, we next measured changes in transcriptional outputs of CD4^+^ T helper cells using bulk RNA sequencing. We found that *Ets1* expression was reduced by ∼30% in *Ets1*-SE^−/−^ Th1 cells but *Fli1* expression remained intact (Fig. 5a). Moreover, the expression of 33 genes was reduced while the expression of 51 genes increased in *Ets1*-SE^−/−^ Th1 cells (Fig. 5b, Table S2). Strikingly, *Ets1*-SE deletion had a specific effect on the Th1 gene expression program, with Th1 signature genes, such as *Ifng* and *Il18r,* selectively and significantly downregulated in *Ets1*-SE^−/−^ Th1 cells compared with wildtype counterparts (Fig. 5c). Consistent with studies showing that the suppression of the Th1 program promotes Th2 differentiation^32^, Th2 signature genes were selectively upregulated in *Ets1*-SE^−/−^ cells polarized under Th1 conditions (Fig. 5d). Unlike the Th1-specific effect of *Ets1*-SE, deletion of this regulatory element did not change the transcriptional landscape of Th2 cells (Fig. 5e). Of note, the effect of *Ets1*-SE deletion on the expression level of *Ets1* and the number of differentially expressed genes was more pronounced in naïve CD4^+^ T cells, implying the role of *Ets1*-SE in the naïve state (Figs. 5a and S5a-c). We postulate that the increase in frequency of connectivity within the multi-enhancer hub across individual alleles after *Ets1*-SE deletion (Fig. 4) is a compensatory mechanism to maintain high expression level of *Ets1*, leading to only a partial reduction in *Ets1* expression. Nonetheless, the partial reduction in *Ets1* expression is sufficient to cause a significant and specific decrease in the Th1-associated gene expression program.

**Figure 5:**
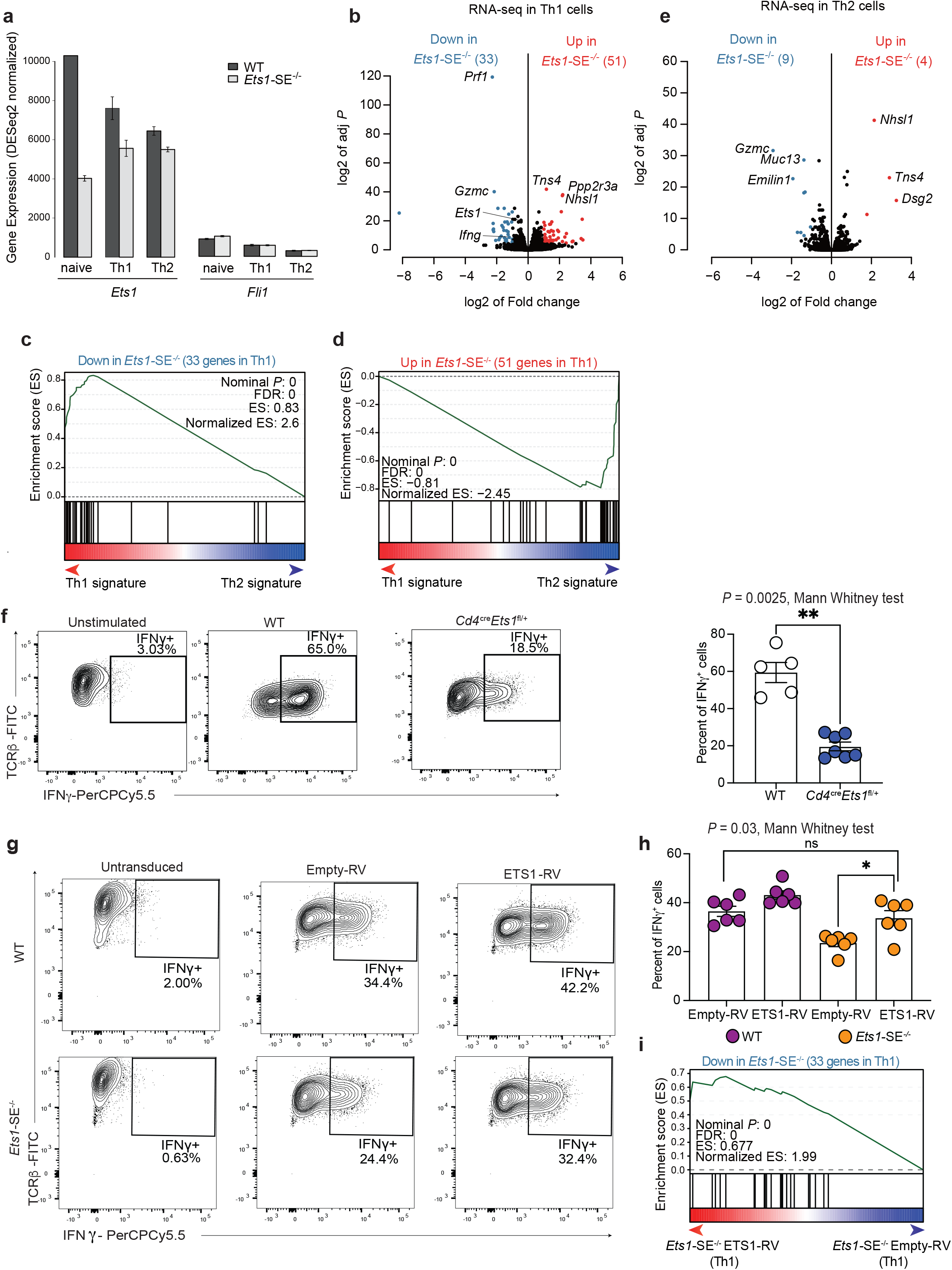
ETS1 dosage controls Th1 differentiation. **a,** Barplot demonstrates normalized mRNA levels of *Ets1* and *Fli1* using bulk RNA-seq experiments in wildtype and *Ets1*-SE^−/−^ naïve CD4^+^ and *in vitro* polarized Th1, and Th2 cells performed in three replicates. **b,e,** Volcano plot demonstrates differential expression analysis of bulk RNA-seq experiments in wildtype and *Ets1*-SE^−/−^ Th1 (b) and Th2 (e) *in vitro* polarized cells studied at day 6. Three replicates were used to perform DESeq2 analysis and |log2FC|>1, adjusted *P* <0.05 were used to determine differentially expressed genes. **c,d,** Pre-ranked Gene Set Enrichment Analysis (GSEA) depicts the enrichment of gene-set for downregulated (c) and upregulated (d) genes in *Ets1*-SE^−/−^ compared with wildtype Th1 cells. The pre-ranked genes were defined based on DESeq2 analysis of wildtype Th1 and Th2 cells. **f,** Flow cytometry results demonstrating frequency of IFNγ expressing cells under unstimulated (left) and *in vitro* Th1 polarization for 6 days in wildtype (middle) and *Ets1* heterozygous mice (*Cd4*^cre^*Ets1*^fl/+^) (right). The right panel shows bar plot of average results from two independent experiments. Each dot represents an individual mouse. Statistical significance was evaluated using non-parametric Mann-Whitney test. ** *P* = 0.0025. **g,h,** Flow cytometry results demonstrating frequency of IFNγ expressing cells when naïve CD4^+^ T cells were retrovirally transduced with empty vector (Empty-RV) or ETS1 expressing vector (ETS1-RV) and were polarized under *in vitro* Th1-differentiating conditions for 6 days. **g**, panel shows representative contour plots, **h**, panel shows bar plot of average results from two independent experiments. Each dot represents an individual mouse. Significance was tested using non-parametric Mann-Whitney test. * *P* = 0.03. **i,** Pre-ranked Gene Set Enrichment Analysis (GSEA) depicts the enrichment of the downregulated genes in *Ets1*-SE^−/−^ compared with wildtype Th1 cells as gene-set. The pre-ranked genes based on DESeq2 analysis of *Ets1*-SE^−/−^ cells transduced with empty-vector and ETS1-expressing vector. Three technical replicates were used for DESeq2 analysis.

### ETS1 dosage controls Th1 differentiation

We next assessed whether partial reduction of the expression level of *Ets1* was responsible for the compromised Th1 differentiation in *Ets1*-SE^−/−^ cells using two complementary approaches. First, we used *Ets1* heterozygous mice (*Cd4*^cre^*Ets1*^fl/+^) that harbor 50% reduction of *Ets1* transcript and protein levels in CD4^+^ T cells. Second, we overexpressed ETS1 protein in *Ets1*-SE^−/−^ Th1 cells that carry 30% less expression of *Ets1* as compared to wildtype cells. Thus, using these *Ets1* dosage experiments, if the *Ets1* expression level controls Th1 differentiation through the *Ets1*-SE element (in *cis*), we expect that cells carrying one copy of the *Ets1* gene, but an intact *Ets1*-SE region, would also fail to differentiate effectively towards the Th1 program. Moreover, we expect that the reconstitution of ETS1 in *Ets1*-SE^−/−^ Th1 cells to the wildtype level can rescue optimal Th1 differentiation. We observed a significant loss of IFNγ expressing cells upon *in vitro* Th1 polarization of CD4^+^ cells from *Cd4*^cre^*Ets1*^fl/+^ mice when compared with wildtype counterparts (Fig. 5f). Furthermore, the exogenous expression of ETS1 into *Ets1*-SE^−/−^ cells using retroviral transduction during *in vitro* Th1 polarization significantly increased the frequency of IFNγ expressing cells compared to empty-vector transduced cells comparable to numbers in wildtype cells (Fig. 5g-h). Transcriptional profiling of *Ets1*-SE^−/−^ Th1 cells with the ectopic expression of ETS1 further confirmed selective rescue of Th1-signature genes (Figs. 5i and S5d,e). Together, our Th1 differentiation assay using *Ets1* heterozygous cells and the rescue experiments in *Ets1*- SE^−/−^ cells implies *Ets1*-SE acts in *cis* at this hyperconnected locus, suggesting that a precise expression level of *Ets1* is required for optimal Th1 differentiation.

### The chromatin accessibility landscape of Th1 cells does not depend on the *Ets1* level

To further dissect the mechanisms of Th1 gene activation, we assessed whether a precise level of *Ets1* expression was required to establish the active enhancer landscape of Th1 cells. We examined the chromatin accessibility landscape of Th1 cells using ATAC-seq and found only a few genomic elements to be differentially accessible between wildtype and *Ets1*-SE^−/−^ Th1 cells (Fig. S6a). Similarly, a small number of genomic regions demonstrated significant alterations in histone acetylation as measured by H3K27ac CUT&RUN^44^ in *Ets1*-SE^−/−^ compared to wildtype Th1 cells (Fig. S6b). Hence, we did not find any strong evidence for the active enhancer landscape of Th1 cells to be dependent on the *Ets1* level. In line with transcriptional profiling, the chromatin accessibility landscapes of wildtype and *Ets1*-SE^−/−^ Th2 cells were virtually indistinguishable (Fig. S6c). Notably, the *Ets1*-SE deletion led to major changes on the chromatin accessibility landscape of naïve T cells (Fig. S6d-i). The comparison of chromatin accessibility landscapes across cell types revealed that the *Ets1*-SE-dependent open chromatin regions in naïve T cells had low level of accessibility in differentiated Th1 cells, suggesting a distinct effect of *Ets1*-SE on the naïve state (Fig. S6g-i). Altogether, the Th1-specific enhancer landscape was largely independent of the *Ets1* level controlled by the *Ets1*-SE element.

### *Ets1* level controls the 3D genome topology of Th1 cells

To assess whether the *Ets1* expression level is required for the spatial localization of enhancers to Th1 signature genes, we created an unbiased map of long-range interactions in CD4^+^ T helper cells using ultra-deep Hi-C^45,46^. Overall, genome organization at the scales of compartments and topologically associating domains (TADs) remained comparable between wildtype and *Ets1*-SE^−/−^ Th1 cells (Fig. S7a-b). However, we observed 1,880 weaker and 325 stronger long-range interactions in *Ets1*-SE^−/−^ as compared to wildtype Th1 cells (Fig. 6a). Genes associated with Th1 biology were located within loop domains with reduced interaction in *Ets1*-SE^−/−^ compared with wildtype cells, suggesting the specific effect of the *Ets1* expression level on optimal Th1-specific genome topology (Fig. 6b). In particular, the long-range interaction at a major loop confining the prototypical Th1 cytokine *Ifng* and its corresponding noncoding-RNA *Tmevpg1* or *NeST*^47,48^ dramatically weakened after *Ets1*-SE deletion (Fig. 6c). Other Th1-associated loci including the cluster of *Il1r-Il18r* genes were misfolded in *Ets1*-SE^−/−^ compared with wildtype Th1 cells (Fig. S8a). Consistent with gene expression and chromatin accessibility measurements, the effect of the *Ets1*-SE deletion on long-range interactions was less pronounced in Th2 cells compared with the effect of this regulatory element on Th1 cells (Fig. S8b). Thus, a precise level of *Ets1* expression controlled by the *Ets1*-SE element is required for the Th1-associated genome topology.

**Figure 6:**
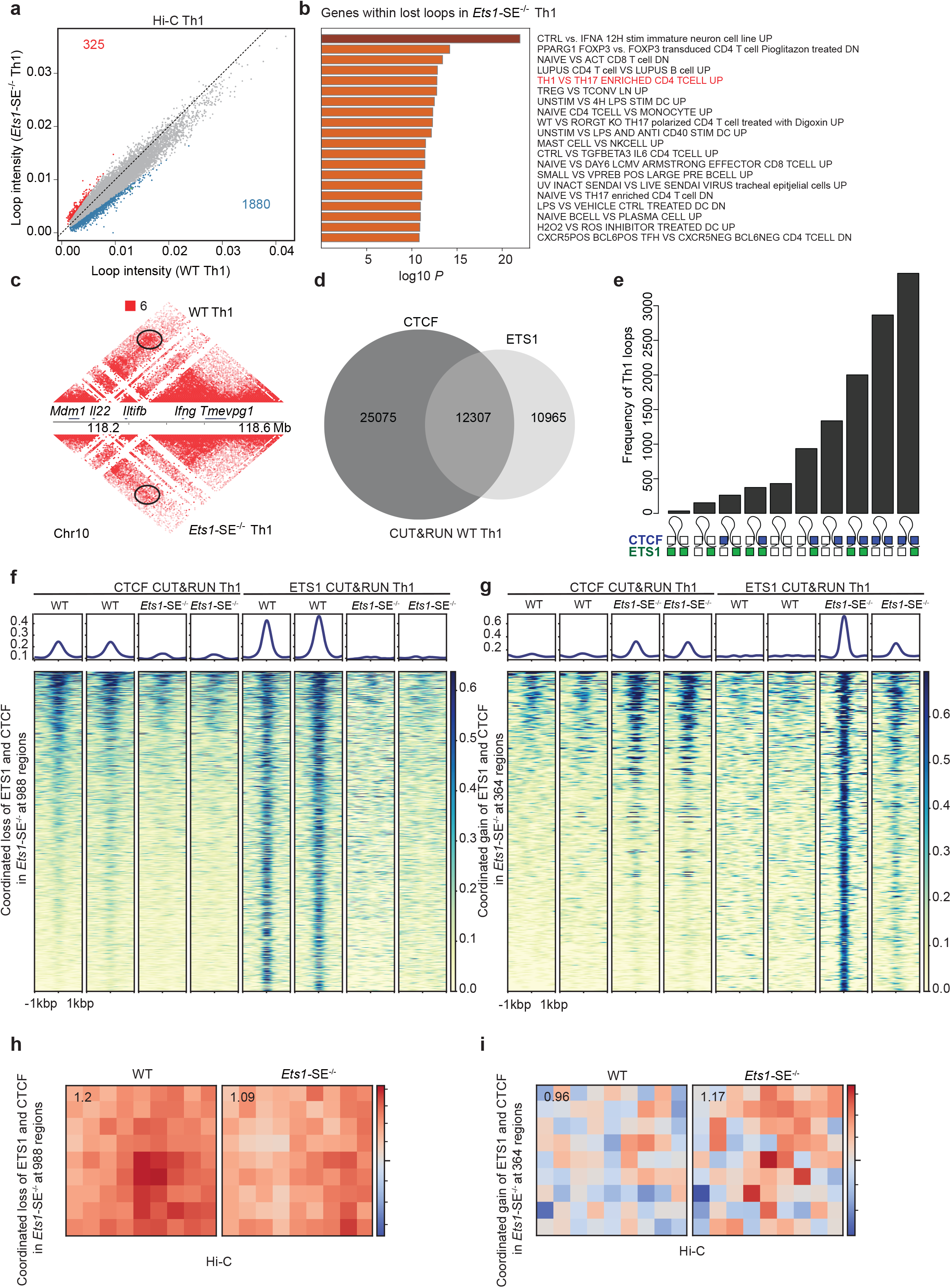
*Ets1* level controls the 3D genome topology of Th1 cells in a CTCF-dependent manner. **a,** Scatter plot depicts intensity of long-range interactions at loops measured by ultra-deep Hi-C between wildtype and *Ets1*-SE^−/−^ Th1 cells. Loops were detected using Mustache^60^ and differential loops were identified using FDR<0.1 & |log2FC|>0.5. Red and blue dots correspond to gained and lost loops in *Ets1*-SE^−/−^ Th1 cells compared with wildtype Th1 cells, respectively. **b,** Barplot depicts *P* values for the enrichment of gene-ontology terms using MetaScape within genes located on *Ets1*-SE dependent loop domains in Th1 cells. Red ontology indicates terms related to Th1 biology. **c,** Heatmap demonstrates ultra-deep Hi-C contact frequencies between wildtype and *Ets1*-SE^−/−^ Th1 cells at the *Ifng* locus. Black circles indicate the differential loop identified in this analysis. The normalized contact frequency corresponding to the strongest pixel is shown in red. **d,** Venn diagram demonstrates the number of genomic regions with co-binding of ETS1 and CTCF measured by CUT&RUN under wildtype Th1 condition. Two replicates were used, and peaks were called by macs2 were combined across different replicates. **e,** UpSet plot demonstrates the number of Th1-associated loops with different binding patterns of CTCF and ETS1 across two loop anchors. Bedtools intersect was used to define overlapping peaks with loop anchors detected in Th1 cells. **f,g** Heatmap demonstrates CTCF and ETS1 occupancy levels at genome regions where both proteins are lost (f) and gained (g) in *Ets1*-SE^−/−^ compared with wildtype Th1 cells. DESeq2 was used to define co-bound regions by ETS1 and CTCF measured by CUT&RUN with differential occupancy between wildtype and *Ets1*-SE^−/−^ Th1 cells using *P*<0.05 & |logFC|>0.5. **h, i**, Loop pileups of long-range interactions anchored at genomic regions described in (f) and (g) with lost (h) and gained (g) occupancy generated by coolpuppy. Hi-C data in wildtype and *Ets1*-SE^−/−^ Th1 cells were used.

### ETS1 and CTCF co-bound sites sensitive to the *Ets1* expression level shape the 3D genome topology of Th1 cells

We next assessed whether partial reduction of *Ets1* could modify the genome-wide occupancy of ETS1 protein and if there was any direct evidence for ETS1 binding affecting Th1-associated long-range interactions. We therefore mapped the global binding pattern of ETS1 protein across biological replicates of Th1 cells using CUT&RUN^44^ (Fig. S8c). Overall, we detected more than 27,000 genomic regions bound by ETS1 in wildtype Th1 cells enriched with the canonical ETS recognition motif and found ∼33% occupancy at promoter regions, suggesting the high quality of our CUT&RUN measurements (Fig. S8d-e, Table S3). The ubiquitously expressed protein CTCF has a prominent role in creating long-range interactions through its convergent binding events, which can block cohesin-mediated loop extrusion^49,50^. Considering the unexpected effect of the level of *Ets1* expression on the 3D genome organization of Th1 cells (Fig. 6a), we next measured the genome-wide occupancy of CTCF using CUT&RUN and found that more than 50% of ETS1 binding events co-occurred at CTCF-bound sites in Th1 cells (Fig. 6d). Moreover, ETS1 and CTCF co-binding was detected at a significant proportion of Th1-associated loop anchors (Fig. 6e), suggesting the potential cooperation of ETS1 and CTCF in establishing 3D genome topology of Th1 cells. To further understand how the reduced expression of *Ets1* weakened long-range interactions in *Ets1*-SE^−/−^ Th1 cells, we compared ETS1 and CTCF occupancies in wildtype and *Ets1*-SE^−/−^ Th1 cells. While the majority of ETS1 and CTCF binding events were not sensitive to the *Ets1* expression level, 998 genomic regions demonstrated a coordinated loss of both ETS1 and CTCF binding in *Ets1*-SE^−/−^ compared to wildtype Th1 cells (Fig. 6f, Table S4). In addition, 364 genomic regions demonstrated a coordinated gain of both proteins in *Ets1*-SE^−/−^ compared to wildtype Th1 cells (Fig. 6g). The subset of ETS1-CTCF co-occupied regions with a coordinated loss of both proteins in *Ets1*-SE^−/−^ Th1 cells demonstrated lower ETS1 and CTCF occupancy compared to the majority of unaffected binding events in wildtype Th1 cells, implying the overall sensitivity of these loci to the *Ets1* expression level (Fig. S8f). Remarkably, the ETS-RUNX cooperative binding motif, which has been characterized as the T cell activation signature of ETS1 binding^21,51^, was enriched at genomic regions which were sensitive to the *Ets1* level (Fig. S8g). STAT2 and Tbox motifs which are recognition sites for Th1-specific transcription factors were also enriched at these ETS1-CTCF lost sites, suggesting the potential cooperation of T-bet and STAT2 transcription factors with ETS1 and RUNX proteins at these loci (Fig. S8g). Linking ETS1-CTCF co-occupied regions to 3D long-range interactions measured by Hi-C, we found that genomic regions whose ETS1 and CTCF co-occupancy were sensitive to the *Ets1* level demonstrated altered long-range interactions in *Ets1*-SE^−/−^ Th1 cells (Fig. 6h,i). These results indicate that a precise level of ETS1 protein was required for CTCF occupancy and the 3D long-range interactions for Th1 regulatory elements. Altogether, the hyperconnectivity of the *Ets1-Fli1* locus controlled the expression level of *Ets1* which was dispensable for the active enhancer signature but required for the Th1-specific DNA folding through recruitment of CTCF. This specific deficit in genome folding due to a partial reduction of *Ets1* led to compromised Th1 differentiation and an allergic response *in vivo*.

## Discussion

Here, we interrogated the functional relevance of a frequently reported 3D structure referred to as a multi-enhancer hub in the context of T cell differentiation. Our systematic prioritization of 3D genome interactions in mouse T cells ranked the *Ets1-Fli1* region as the second most hyperconnected locus after the *Bcl11b* region with an unusual degree of enhancer connectivity. This multi-enhancer locus was conserved in human T cells and was found to be a hotspot for SNPs associated with allergic diseases. To better understand the functional relevance of the hyperconnectivity of this genetic hotspot in T cell biology, we generated a new mouse strain and deleted a noncoding regulatory element homologous to the allergy-associated polymorphic region in the human genome. While T cell development remained intact, Th1 differentiation was compromised after the genetic deletion of this regulatory element. Detailed mechanistic investigation demonstrated the link between the hyperconnectivity of the *Ets1-Fli1* locus*, Ets1* expression level, CTCF recruitment, and long-range interactions required for the Th1 gene expression program. Although it has been shown that the complete ablation of *Ets1* can lead to changes in CTCF recruitment^52^, our study is the first demonstration of the sensitivity of genome organization to the *Ets1* expression level as a mechanism for predisposition to immune-mediated diseases. Related to our findings, it has been known that the graded expression of interferon regulatory factor-4 (IRF-4) coordinates isotype switching with plasma cell differentiation^53^. We speculate that IRF-4 or other transcription factors may follow a similar dose-dependent mechanism and reorganize the genome by interacting with CTCF at specific genome regions.

Since the completion of the Human Genome Project, research on the genetic basis of Mendelian traits has been extremely successful. In most cases, the rare single-gene disorders masquerade a multifactorial trait in their clinical phenotype^54^, but detailed clinical examination and studying ‘human allelic knockouts’ have guided to one gene responsible for the disease in those particular families. Thus, genetic ablation of an individual gene in rodents became a powerful tool to dissect the molecular processes of such monogenic diseases. Despite the success of genetic strategies in Mendelian traits, human genetics has been less successful in dissecting complex conditions which are in fact more commonly found across populations. Unlike Mendelian traits that can be modeled as gene knockouts, complex traits might be caused by single-nucleotide variants disrupting transcriptional enhancers. However, the link between sequence variation, cellular context, 3D genome folding, and the extent of change in gene expression in most complex diseases remains largely unknown. Here we presented a mouse model, which was inspired by the mathematical analysis of genome organization data, reporting how noncoding elements can control the precise dosage of a key transcription factor through the formation of a multi-enhancer hub. It is worth mentioning that the heterozygous alleles of transcription factors such as HNF1A^55^ or FoxA2^56^ can impact target genes, leading to developmental defects. Yet, our study highlights dosage control from the perspective of regulatory elements of the transcription factor gene, not the coding region of the transcription factor gene itself.

Our optical probing using the Oligopaint technology revealed an unexpected reorganization of the multi-enhancer hub in the absence of a key regulatory element. We therefore expect that genome variation might also be able to modify gene-enhancer interactions in a similar fashion. As of today, we are limited in these experiments to select three genomic loci, but chromatin tracing experiments^57^ for the entire multi-enhancer hub could allow us to better understand the dynamics of multi-enhancer interactions at the single-cell resolution. Considering that individuals carrying risk factors for genetic predispositions to common diseases are far from gene-knockout mouse models, we reason that comprehensive strategies following the integrative approaches used in this study can shed light on molecular mechanisms through which single-nucleotide variants can affect the gene expression programs sometimes in subtle ways which can lead to substantial clinical phenotypes.

---

**Figure S1:**
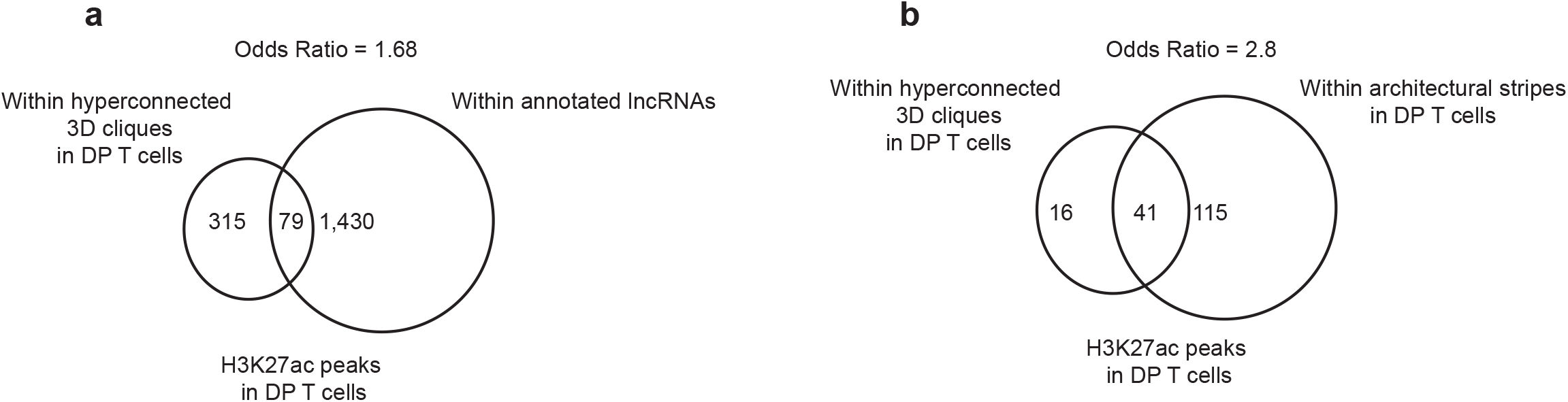
Colocalization of multi-enhancer hubs with ncRNAs and architectural stripes. **a,** Venn diagram depicts the number of overlaps in genomic regions which scored as hyperconnected multi-enhancer hubs in DP T cells and annotated noncoding (nc)RNAs. Annotation of ncRNA was performed using gencode.vM10.annotation.gtf file. Odds ratio was calculated by creating contingency tables of overlapping genomic regions. **b**, Venn diagram depicts the number of overlaps in genomic regions which scored as hyperconnected multi-enhancer hubs in DP T cells and architectural stripes detected in DP T cells using Stripenn^14^. Odds ratio was calculated by creating contingency tables of overlapping genomic regions.

**Figure S2:**
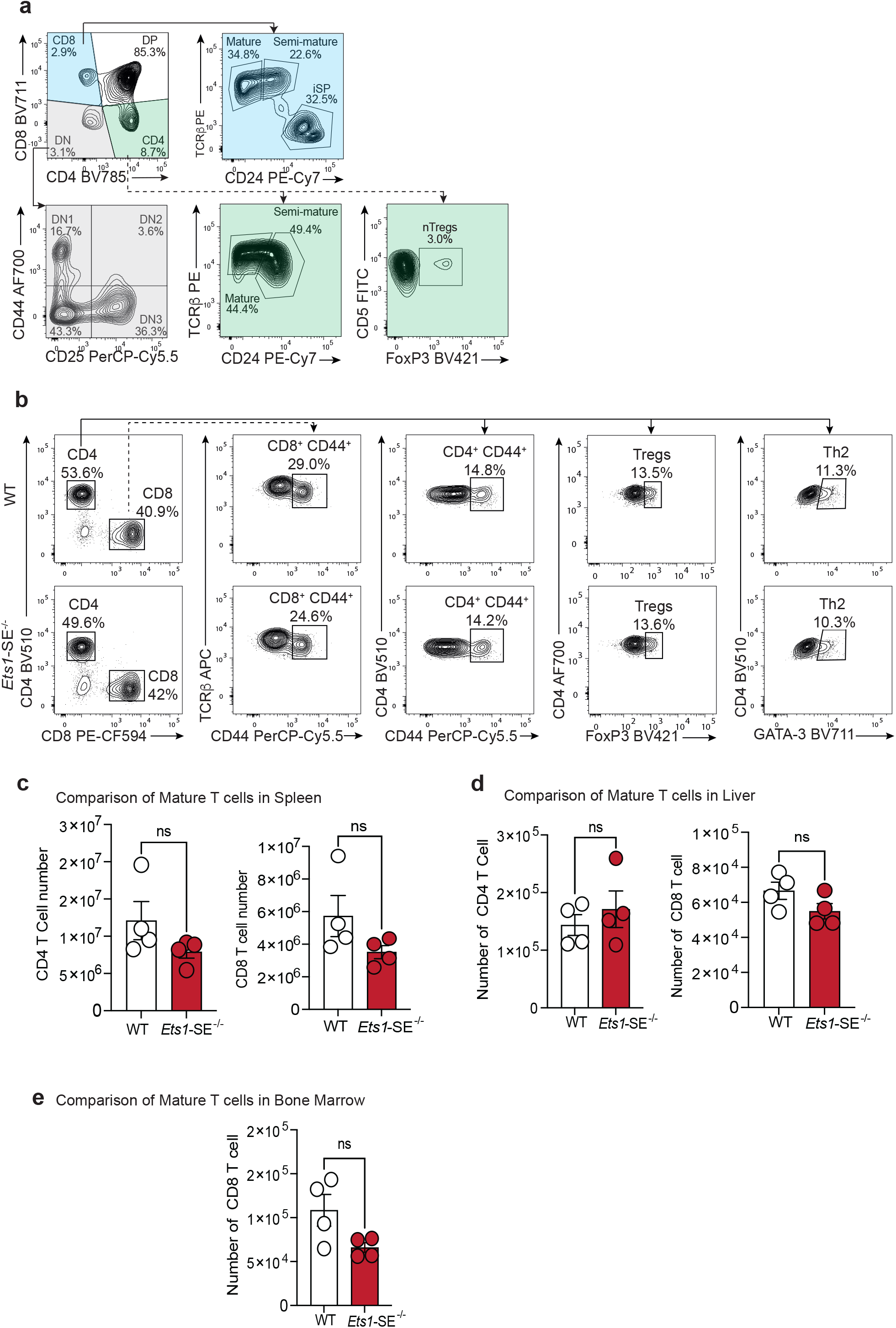
*Ets1*-SE is dispensable for thymic T cell development and periphery colonization. **a,** Gating strategy used to identify thymic T cell population of wildtype or *Ets1-SE*^−/−^ related to **Figure. 2a**. Cells were pre-gated on SSC-A/FSC-A, Singlets and Live (Viability^−^) cells. Colored boxes indicate sequential gating. **b,** Gating strategy used to identify lung parenchyma immune cell populations of wildtype or *Ets1-* SE^−/−^ at steady state related to Figure 2b. Cells were pre-gated on SSC-A/FSC-A, Singlets and Live (Viability^−^) cells. **c,** Quantification of mature CD4^+^ and CD8^+^ T cells in the spleen of wildtype or *Ets1-*SE^−/−^ animals at steady state. Cells were pre-gated on SSC-A/FSC-A, Singlets and Live (Viability^−^) cells. Data are representative of one experiment. Each dot represents an individual mouse (wildtype n= 4 and *Ets1-*SE^−/−^ n=4). Error bars = SEM; and *P*: ns = not significant (Mann-Whitney U test). **d,** Quantification of mature liver CD4^+^ and CD8^+^ T cells of wildtype or *Ets1-*SE^−/−^ animals at steady state. Cells were pre-gated on SSC-A/FSC-A, Singlets and Live (Viability^−^) cells. Data are representative of one experiment. Each dot represents an individual mouse (wildtype n= 4 and *Ets1-*SE^−/−^ n=4). Error bars = SEM; and *P*: ns = not significant (Mann-Whitney U test). **e,** Quantification of mature bone marrow (BM) CD8^+^ T cells of wildtype or *Ets1-*SE^−/−^ animals at steady state. Cells were pre-gated on SSC-A/FSC-A, Singlets and Live (Viability^−^) cells. Data are representative of one experiment. Each dot represents an individual mouse (wildtype n= 4 and *Ets1-*SE^−/−^ n=4). Error bars = SEM; and *P*: ns = not significant (Mann-Whitney U test).

**Figure S3:**
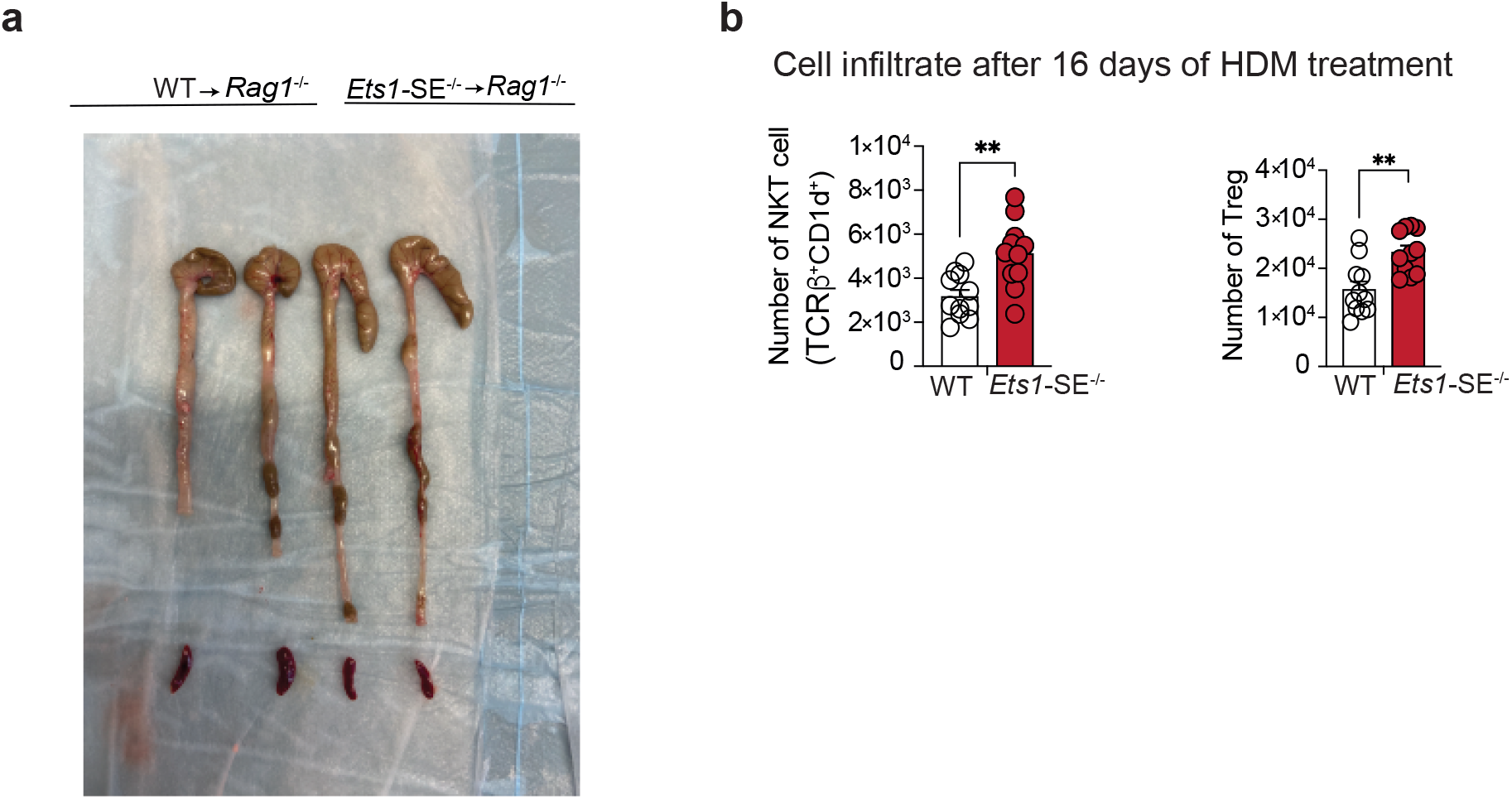
*Ets1*-SE deletion limits Th1-mediated inflammation and enhances Th2 responses. **a,** Picture of colon in *Rag1*^−/−^ mice that received either 1×10^6^ FACS sorted TCRβ^+^, CD4^+^, CD45RB^High^ naïve wildtype or *Ets1-*SE^−/−^ CD4^+^ T cells 6 weeks post transfer. **b,** Quantification of lung parenchyma NKT cells (TCRβ^+^, CD1d^+^) and Tregs (TCRβ^+^, CD4^+^, FoxP3^+^) of wildtype or *Ets1-*SE^−/−^ animals 16 day after the initial HDM challenge. Cells were pre-gated on SSC-A/FSC-A, Singlets and Live (Viability^−^) cells. Data are a pool of two independent experiments. Each dot represents an individual mouse (wildtype n= 11 and *Ets1-*SE^−/−^ n=11). Error bars = SEM; and ** *P*: <=0.01 (Mann-Whitney U two-tailed test).

**Figure S4.**
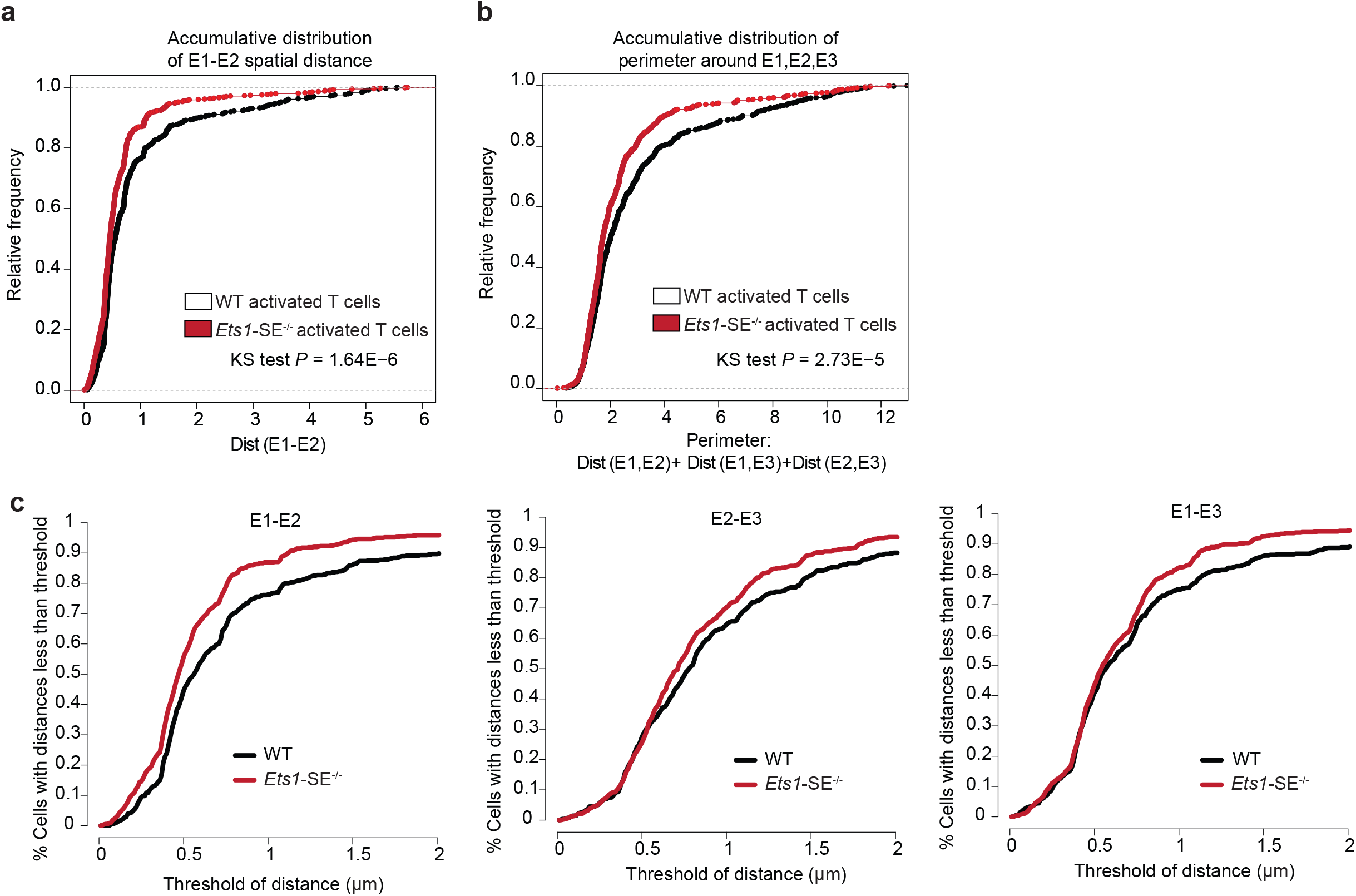
Genome topology of the *Ets1* locus changes after *Ets1*-SE deletion. **a**,**b** Accumulative distribution demonstrating the spatial distance formed between E1 and E2 probes (a) and the perimeter formed by E1, E2, and E3 (b) in CD4^+^ T cells 18 hours after activation with T cell receptor signaling per genotype (KS: Kolmogorov-Smirnov test & MW: Mann-Whitney test). More than 500 CD4^+^ cells were imaged using widefield microscopy from one wildtype and one *Ets1*-SE^−/−^ mouse. **c**, Accumulative distribution demonstrating percentage of activated CD4^+^ T cells where spatial distance between genomic regions corresponding to E1 and E2, E2 and E3, and E1 and E3 probes are less than a range of threshold (x-axis).

**Figure S5:**
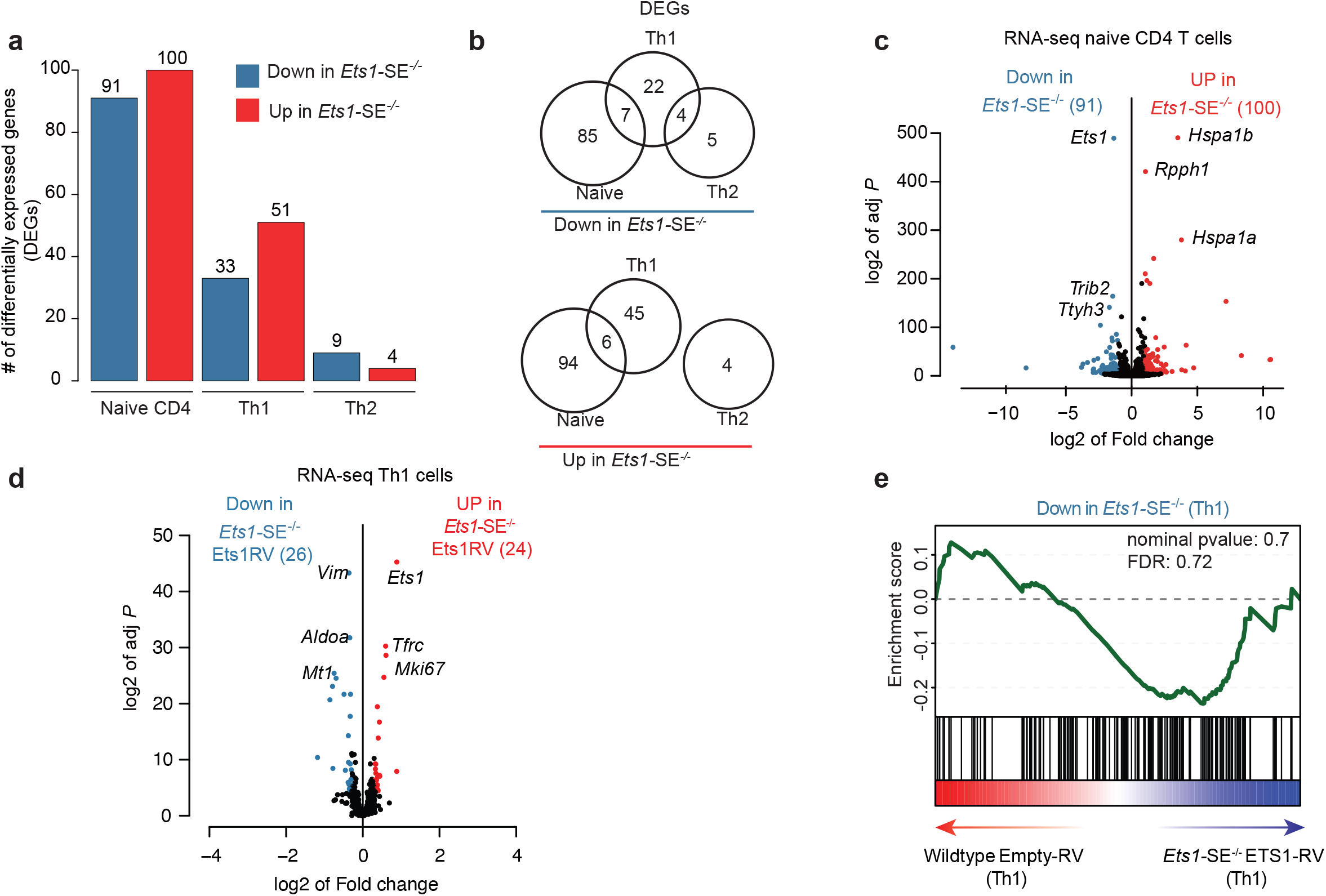
Transcriptional outputs of Th1 cells depend on *Ets1*-SE. **a,** Barplot depicts number of differentially expressed genes detected by DESeq2 using bulk RNA-seq measurements in wildtype and *Ets1*-SE^−/−^ cells under naïve, Th1 and Th2 conditions. Three replicates were used to perform DESeq2 analysis and |log2FC|>1, adjusted *P* <0.05 were used to determine differentially expressed genes. **b,** Venn diagram depicts the number of shared and unique genes differentially expressed in wildtype and *Ets1*-SE^−/−^ cells under naïve, Th1 and Th2 conditions. Three replicates were used to perform DESeq2 analysis and |log2FC|>1, adjusted *P* <0.05 were used to determine differentially expressed genes. **c,** Volcano plot demonstrates differential expression analysis of bulk RNA-seq experiments in wildtype and *Ets1*-SE^−/−^ naïve T cells. Three replicates were used to perform DESeq2 analysis and |log2FC|>1, adjusted *P* <0.05 were used to determine differentially expressed genes. **d,** Volcano plot demonstrates differential expression analysis of bulk RNA-seq experiments in *Ets1*-SE^−/−^ Th1 cells transduced with empty or ETS1 retroviral vector. Three replicates were used to perform DESeq2 analysis and |log2FC|>0.5, adjusted *P* <0.05 were used to determine differentially expressed genes. **e,** Pre-ranked Gene Set Enrichment Analysis (GSEA) depicts the enrichment using gene-set for downregulated genes in *Ets1*-SE^−/−^ compared with wildtype Th1 cells. The pre-ranked genes are based on DESeq2 analysis of *Ets1*-SE^−/−^ cells transduced with ETS1-expressing vector compared with wildtype cells transduced with empty vector.

**Figure S6:**
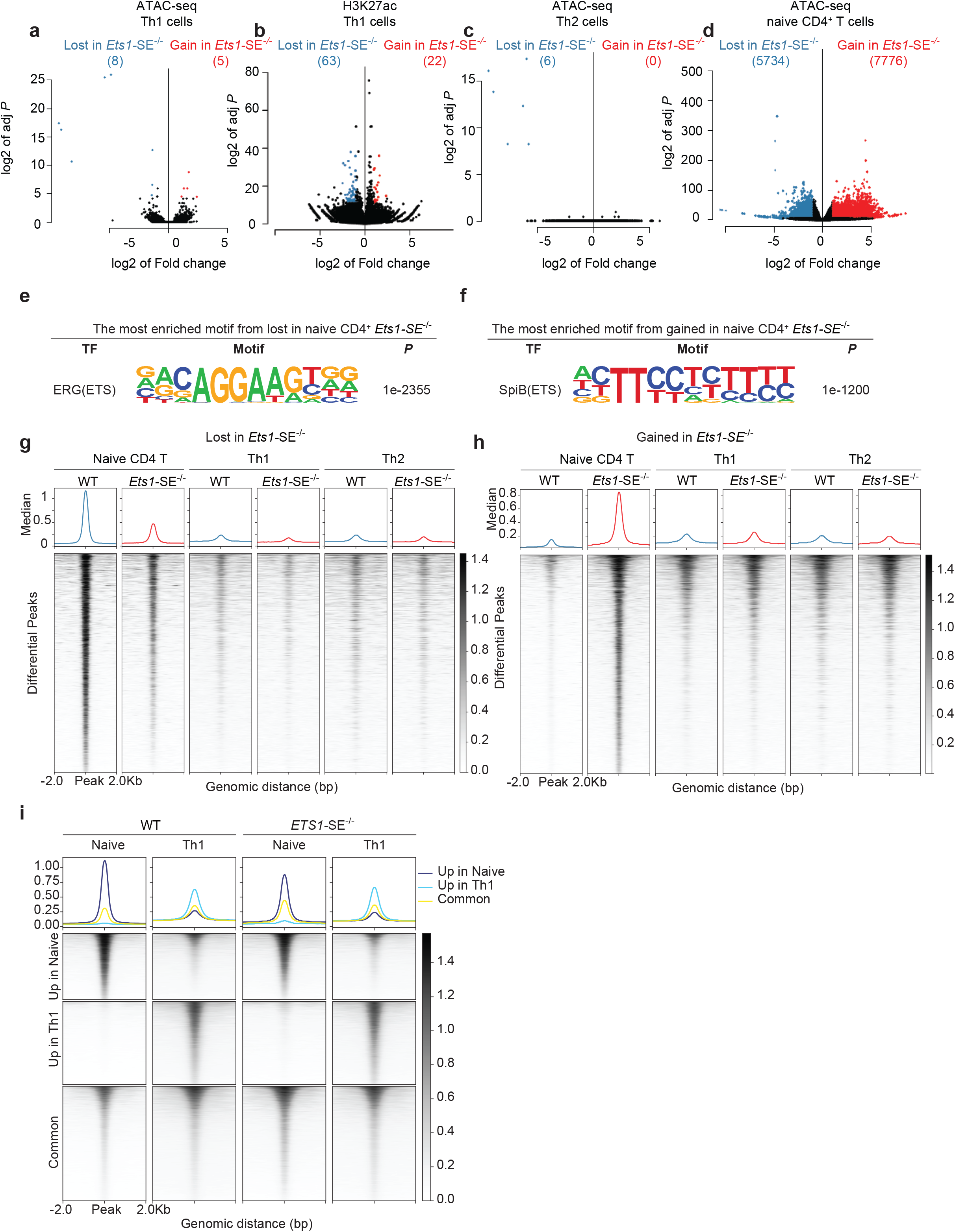
Chromatin accessibility landscape of Th1 cells does not depend on the *Ets1* level. **a,** Volcano plot demonstrates differential accessibility analysis of ATAC-seq experiments in wildtype and *Ets1*-SE^−/−^ CD4^+^ Th1 cells. Three replicates were used to perform DESeq2 analysis and |log2FC|>1, adjusted *P* <0.05 were used to determine differentially accessible regions. **b,** Volcano plot demonstrates differential H3K27ac analysis of CUT&RUN experiments in wildtype and *Ets1*-SE^−/−^ Th1 cells. Three replicates were used to perform DESeq2 analysis and |log2FC|>1, adjusted *P* <0.05 were used to determine differentially acetylated regions. **c,** Volcano plot demonstrates differential accessibility analysis of ATAC-seq experiments in wildtype and *Ets1*-SE^−/−^ Th2 cells. Three replicates were used to perform DESeq2 analysis and |log2FC|>1, adjusted *P* <0.05 were used to determine differentially accessible regions. **d,** Volcano plot demonstrates differential accessibility analysis of ATAC-seq experiments in wildtype and *Ets1*-SE^−/−^ naïve CD4^+^ T cells. Three replicates were used to perform DESeq2 analysis and |log2FC|>1, adjusted *P* <0.05 were used to determine differentially accessible regions. **e,f** Seqlogo depicts *de novo* motif analysis using homer in lost (e) and gained (f) accessible chromatin regions in wildtype and *Ets1*-SE^−/−^ naïve CD4^+^ T cells. **g,h** Heatmap demonstrates the intensity of accessible chromatin in differential regions of naïve T cells. Lost (g) and gained (h) peaks are demonstrated. **i**, Heatmap demonstrates intensity of accessible chromatin in differential regions between naïve T cells and Th1 cells.

**Figure S7:**
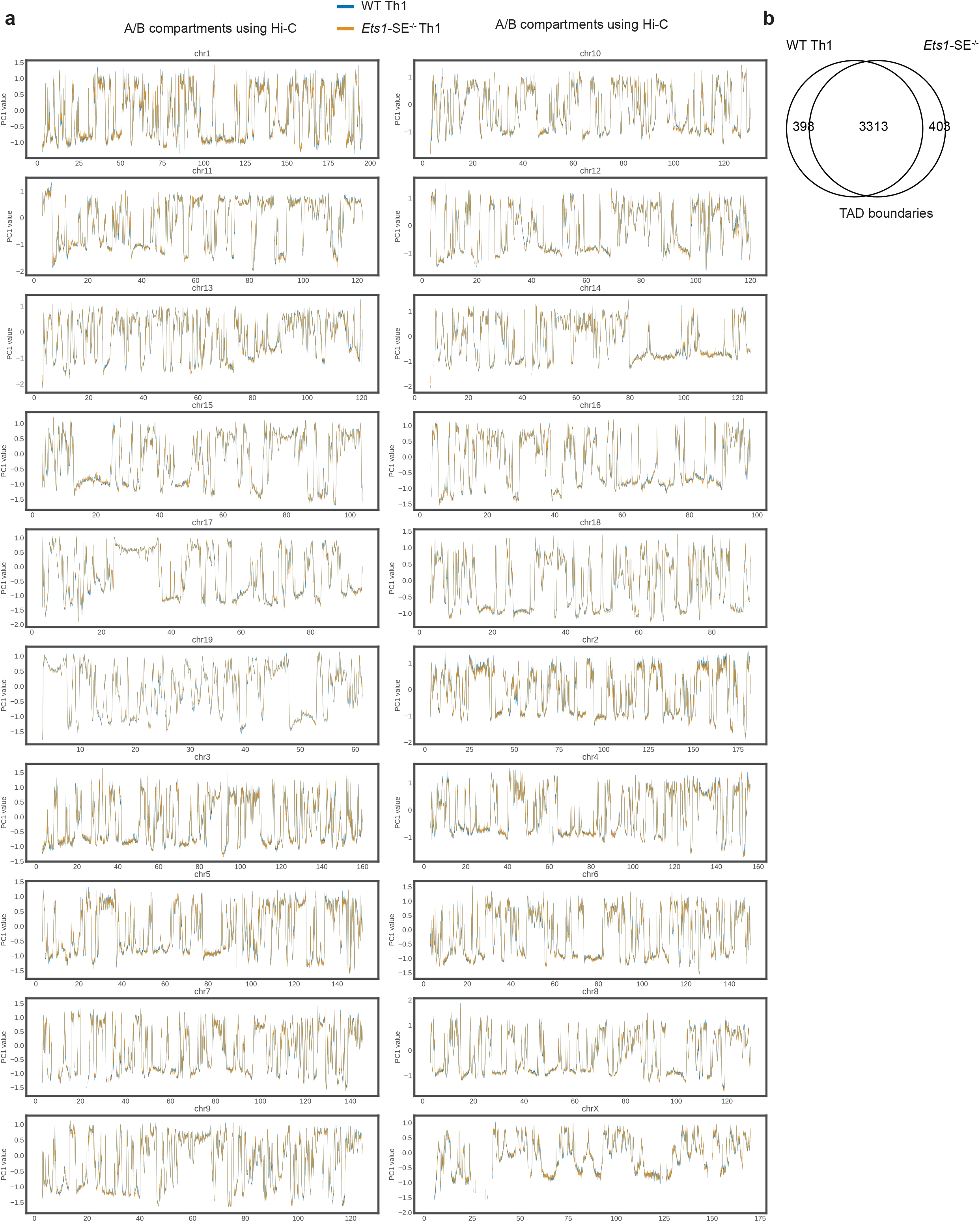
Compartment and TAD analysis in wildtype and *Ets1*-SE^−/−^ Th1 cells. **a,** Plots demonstrate compartment analysis using PC1 values in wildtype and *Ets1*-SE^−/−^ Th1 cells using Hi-C across 20 chromosomes in mice. **b,** Venn diagram demonstrates shared and overlapping boundaries detected by insulation score analysis in wildtype and *Ets1*-SE^−/−^ CD4^+^ Th1 cells using Hi-C.

**Figure S8:**
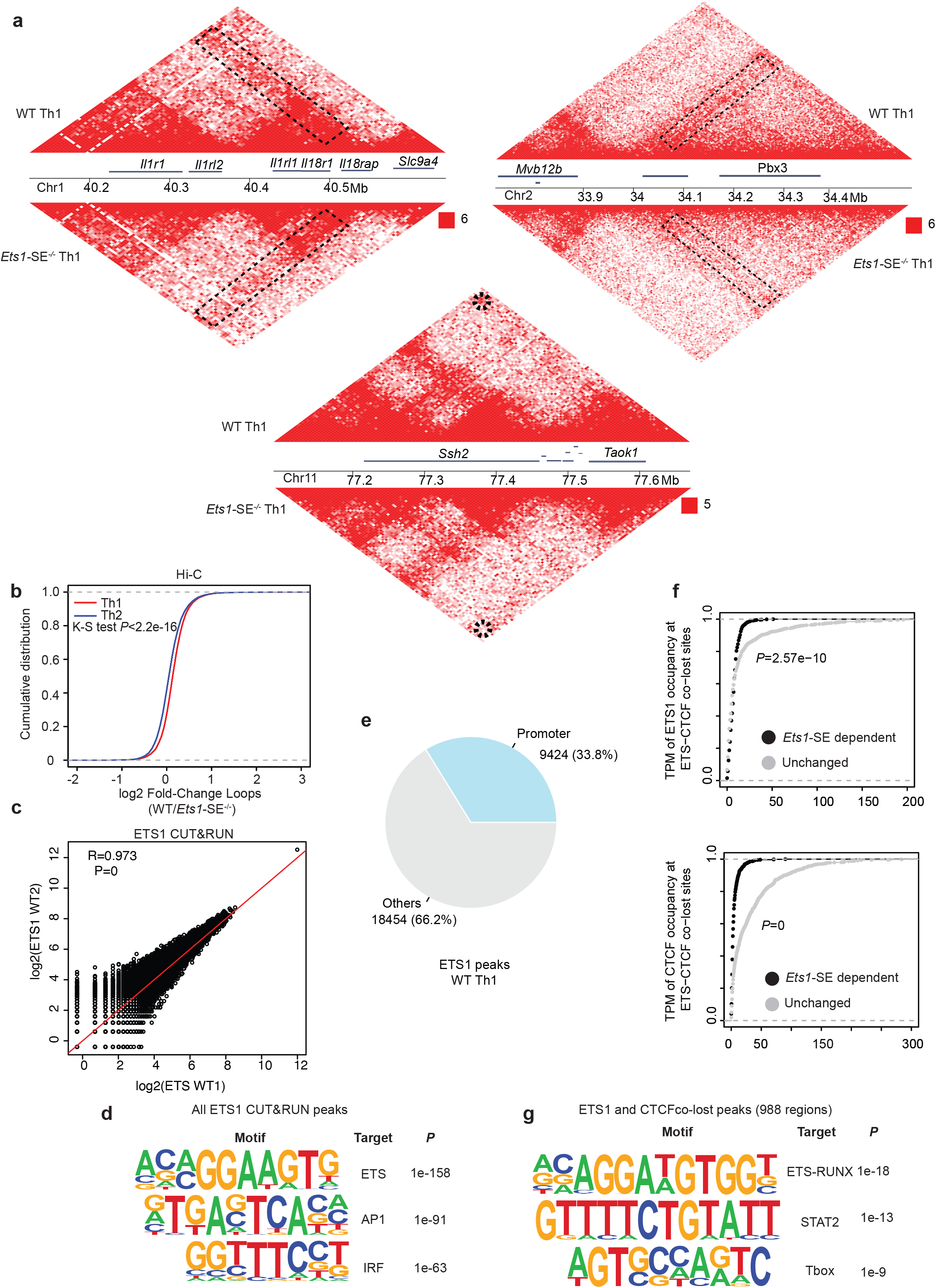
*Ets1* level controls the 3D genome topology of Th1 cells in a CTCF-dependent manner. **a,** Heatmap demonstrates ultra-deep Hi-C contact frequencies between wildtype and *Ets1*-SE^−/−^ Th1 cells at three representative loci including the *Il18r* locus. Black circles indicate the differential loop identified in this analysis. The normalized contact frequency corresponding to the strongest pixel is shown in red. **b,** Accumulative plot demonstrates log2FC for loop intensity measured by Hi-C between wildtype and *Ets1*-SE^−/−^ cells polarized under Th1 and Th2 conditions. (KS: Kolmogorov-Smirnov test) **c,** Scatter plot demonstrates correlation of ETS1 intensity in biological replicates of CUT&RUN. At least two replicates were generated for this experiment. **d,** Seqlogo depicts *de novo* motif analysis using homer in ETS1 CUT&RUN measurements in wildtype Th1 cells. **e**, Pie-chart depicts the percentage of promoter overlapping ETS1 binding events in ETS1 CUT&RUN measurements in wildtype Th1 cells. **f**, Accumulative distribution compares the occupancy of ETS1 (or CTCF) at ETS1-CTCF co-occupied sites which are lost in *Ets1*-SE^−/−^ Th1 cells and the majority of ETS1-CTCF co-occupied sites that are not sensitive to the *Ets1* level. **g**, Seqlogo depicts *de novo* motif analysis using homer at ETS1-CTCF co-occupied sites (defined in Fig. 6f) which are lost in *Ets1*-SE^−/−^ Th1 cells and are sensitive to the *Ets1* level.

## Materials availability

This study did not generate new unique reagents.

### Data and code availability

The accession number for the HiC, CUT&RUN, ATAC-seq and RNA-seq reported in this study is NCBI GEO: GSE211178.

## Methods

### EXPERIMENTAL MODEL AND SUBJECT DETAILS

#### Mice

The *Ets1*-SE^−/−^ mice were generated using the CRISPR/Cas9 system as previously described^61^. The sgRNA sequences 5’ GGACGTTGTGCACCTAGGATTGG and 3’ ATAAACGTCAATAATGGTATAGG were used for the generation of knockout mice and allowed the deletion of the following DNA regions referred to as *Ets1*-SE: chr9:32,928,966-32,904,069 (mm10 genome). *Ets1*-SE^−/−^ mice were backcrossed onto the C57BL/6 background for at least three generations to control potential off-target effects. *Rag1*^−/−^ mice (B6.129S7-Rag1tm1Mom/J; Strain #:002216) were purchased from the Jackson Laboratory. CD4^cre^Ets1^fl/fl^ mice were generated by Dr. Barbara Kee’s laboratory at the University of Chicago by insertion of loxP sites flanking exons 7 and 8 of the *Ets1* locus. Ets1^fl/+^ mice were then crossed to mice expressing the Cre recombinase under the *Cd4* promoter (CD4^cre^) to generate *Cd4*^cre^ *Ets1*^fl/+^ mice. Spleens and lymph nodes (LNs) from these mice were used for *in vitro* polarization assays. All mice were bred and maintained under pathogen-free conditions at an American Association for the Accreditation of Laboratory Animal Care accredited animal facility at the University of Pennsylvania. Mice were housed in accordance with the procedures outlined in the Guide for the Care and Use of Laboratory Animals under an animal study proposal approved by an institutional Animal Care and Use Committee. All experiments were performed using 6- to 12-weeks old age and sex matched mice and using both males and females.

#### Tissue and cell Preparation

Different organs were extracted from the mice for T cell phenotyping study, including thymus, lungs, liver, spleen, lymph nodes and bone marrow. All red blood cells were lysed using ACK lysis buffer (GIBCO). Single-cell suspensions from spleen, thymus and lymph nodes were isolated by physical dissociation of tissues and filtered through a 70μm cell strainer (Falcon) in RPMI-1640 media (Invitrogen) containing 1% fetal bovine serum (FBS) (Fisher Scientific). For bone marrow cell isolation, femurs were collected and crushed using a mortar and pestle. Lungs were isolated, minced with scissors, and digested in PBS containing FBS (2%), Collagenase D (1mg/ml), DNase I (0.2mg/ml) for 35 minutes at 37°C with shaking at 200 RPM. The digested lungs were then passed through a 70μm cell strainer. Mice were perfused with 10ml PBS and then transferred to DMEM on ice. The liver was then removed from media and mechanically dissociated using a tissue grinder, then filtered through a 100μm cell strainer. To pellet hepatocytes, the digested livers were centrifuged at 20g for 5min at 4°C. Leukocytes were then purified over an 80/40% Percoll gradient.

#### CD4^+^ T cell isolation and polarization

Splenocytes were isolated from mouse spleen and lymph nodes. Spleen cells were subjected to ACK lysis buffer (Gibco, Invitrogen) to remove red blood cells. Naïve CD4^+^ T cells were enriched using negative selection beads (STEMCELL, Cat# 19765) following manufacturer’s recommendations. Purity of naïve cells were assessed using flow cytometry and found to be >=90% pure. Cells were cultured in RPMI 1640 medium (Invitrogen), supplemented with 10% fetal bovine serum (Fisher Scientific), 1mM sodium pyruvate (Gibco), 1% non-essential amino acids (Gibco), 1X GlutaMAX (Gibco), 1% HEPES (Gibco), 1% penicillin-streptomycin and 0.1% 2-Mercaptoethanol (Gibco). For *in vitro* polarization, flat bottom 96-well plates were coated with 2 μg/mL of anti-CD3 (Clone: 145-2C11, BioLegend, Cat#100302) in PBS overnight at 4°C or 4 hrs at 37°C. For Th1 polarization, 0.2×10^6^ naïve CD4^+^ T cells were cultured in presence of 10 ng/mL of recombinant IL-12 (BioLegend, Cat#577002), 1 ng/mL of recombinant IL-2 (BioLegend, Cat#575402) and 1 μg /mL of soluble anti-CD28 (Clone: 37.51, BD Biosciences, Cat# 553294) for 6 days at 37°C. Th2 polarization was induced by cultivating 0.2×10^6^ naïve CD4^+^ T cells in complete RPMI supplemented with 10 μg /mL of anti-IFNγ (Clone: XMG1.2, BioXCell, Cat#BE0055), 50 ng/mL of recombinant IL-4 (BioLegend, Cat# 574302) and 1 μg/mL of soluble anti-CD28 for 6 days at 37°C. For Th17 polarization, 0.2×10^6^ naïve CD4^+^ T cells were cultured in complete RPMI containing 10 μg/mL of anti-IFNγ, 10 μg/mL of anti-IL-4 (Clone: 11B11, BioXCell, Cat# BE0045), 2 ng/mL of recombinant TGFβ (BioLegend, Cat#763102), and 20 ng/mL of recombinant IL-6 (BioLegend, Cat# 575702) and 1 μg/mL of soluble anti-CD28 for 6 days at 37°C. At day 3 post stimulation, half of the medium was removed and replaced by fresh media containing 2X cytokines concentration of Th1, Th2 or Th17-related polarizing medium.

#### ETS-1 retroviral transduction experiments

cDNA encoding the short isoform of ETS-1 (p54) was cloned into pENTR vector and then into the destination vector MSCV-IRES-Thy1.1 DEST (Addgene: 17442) using Gateway cloning strategy (Gateway Clonase II, Invitrogen). To generate retroviral particles for Ets1-overexpression, 293T cells were purchased from ATCC. Briefly, HEK- 293T cells were maintained in high glucose DMEM medium 1X with L-Glutamine (Invitrogen), supplemented with 100 U/mL penicillin and 100 mg/mL streptomycin (Gibco) with 10% FBS. Cells were maintained at low passage number (< 12), at 70-80% confluency, and were grown at 37°C and 5% CO2. Retroviral vectors were packaged in HEK 293T cells. Briefly, 4×10^6^ HEK 293T cells were plated in 5 ml media in 10 cm dishes on the day prior to transfection. During transfection, 15 μg of MSCV-Thy1.1-Ets1 plasmid was co-transfected with packaging plasmid, 15 μg of pCL-Eco, using Lipofectamine 3000 (Invitrogen). The MSCV-Thy1.1-EV (Empty vector) plasmid was also similarly transfected. The cells were returned to the incubator for 6 hours. Subsequently, the medium was changed to fresh media. Virions were collected 24 and 48 hrs after transfection, snap-frozen, and stored at −80°C for future use.

#### CD4^+^ T cell transduction

Primary naïve CD4^+^ T cells were transduced by addition of virions to culture media supplemented with polybrene (Sigma-Aldrich, cat# H9268) at 4 μg/mL final concentration, followed by centrifugation at 32°C for 1 hour at 3000 rpm. Cells were returned to incubator at 37°C for at least 4 hrs. Subsequently, the medium was changed to fresh culture media supplemented with Th1 polarizing cytokines. Transduction efficiency was checked next day and frequency of polarizing cells was checked on day 6 after transduction.

#### Antibodies, Flow cytometry, and cell sorting

All antibodies were diluted in FACS buffer (PBS + 2% FBS + 2mM EDTA) and used to stain single cell suspensions for 30 minutes at 4°C. First, dead cells were stained by incubation of cell suspension in Viability Dye (eFluor780, eFluor506, or Aqua) diluted in PBS for 10mins at 4°C. Then after a wash with PBS, cells were staining with surface antibodies diluted in FACS buffer (PBS + 2% FBS + 2mM EDTA) and washed with FACS buffer. Cells were then either fixed for intracellular staining using the Foxp3 staining buffer (eBioscience) or were fixed with 2% PFA. For intracellular staining, fixed cells were washed with permwash buffer (eBioscience) and incubated with intracellular antibodies diluted in permwash buffer for 30 mins or left overnight. Cells were washed with permwash buffer and resuspended in FACS buffer with the addition of 123count eBeads (ThermoFischer Scientific, ref:01-1234-42) following manufacturer’s recommendations for cell counting. For cell sorting, cells were stained at a concentration of 100×10^6^ cells/ml in FACS buffer (PBS + 2% FBS + 2mM EDTA) for 30 mins at 4°C then filtered through a 70μm filter prior acquisition and sort using either a 100μm or 70μm nozzle on a BD FACS Aria II SORP under aseptic conditions. Cells were sorted in FACS buffer containing 10% of FBS.

### Flow Cytometry on *in vitro* polarized cells

For flow cytometry, *in vitro* polarized cells were stimulated with PMA, Ionomycin and Golgi Plug for 4 hrs at 37C. Subsequently, they were stained with a viability dye (1:500 L/D Aqua), and extracellular stains with anti-mouse CD4-APC (1:400, clone RM4-5, BioLegend Cat#100516), anti-mouse TCRb chain-FITC (1:400, clone H57-597, BD Biosciences Cat#553170). Cell were then fixed using Foxp3/ Transcription factor Staining buffer Set (eBioscience, Thermo Cat# 00-5523-00) followed by intra-cellular stains with anti-mouse IFNγ-PerCP/Cyanine5.5 (1:200, clone XMG1.2, BioLegend Cat#505821), T-bet monoclonal antibody-PE-Cyanine 7 (1:200, clone 4B10, eBioscience, Cat# 25-5825-80), anti-mouse IL-13-PE eFluor 610 (clone eBio13A, Fisher Cat# 61-7133-80), anti-GATA3 BV711 (BD Biosciences Cat# 565449), anti-mouse IL-17 PE (clone TC11-18H10.1, BioLegend, Cat# 506903), anti-mouse RORgt BV421 (BD Biosciences Cat#562894). Cells were washed and resuspended in PBS for flow cytometry. Data were collected on an LSR II running DIVA software (BD Biosciences) and were analysed using FlowJo software v10.8.1.

#### *In vivo* House Dust Mite (HDM) extract exposure model

HDM extracts (*Dermatophagoides pteronyssinus* extracts; Greer Laboratories lot #361863, 385930) were used to induce allergic airway inflammation and protocol was adapted following^62^. Briefly, mice were sensitized intranasally with 20 µg HDM extracts on day 0 and subsequently challenged intranasally with 10 µg/mouse per day on days 7–13. Three days after the last challenge, mice were anesthetized and used either for immune cell quantification in the lung parenchyma or *ex vivo* restimulation. For lung parenchyma experiments, lungs were isolated, minced with scissors, and digested in PBS containing FBS (2%), Collagenase D (1mg/ml), DNase I (0.2mg/ml) for 35 minutes at 37°C with shaking at 200 RPM. The digested lungs were then passed through a 70μm cell strainer. Cells were resuspended in PBS then split in order to perform lung parenchyma cell infiltration or *ex vivo* restimulation. For *ex vivo* restimulation, cells were plated into round-bottom 96 well plates and stimulated in complete RPMI media containing PMA (Sigma; final concentration 100ng/mL) and Ionomycin (Sigma; final concentration 10ng/mL) and GolgiPlug (1X - BD Bioscience). Cytokines and transcription factor expression were measured by intracellular staining using the “Foxp3 staining buffer” (Ebioscience).

#### Induction and Evaluation of colitis

CD4^+^ T cells were enriched from spleen and lymph node cell suspensions using a cocktail of biotinylated antibodies containing anti-CD8 (53-6.7), anti-CD19 (6D5), anti-BB20 (RA3-6B2), anti-Gr1 (RB6-8C5), anti-TCRγ8 (GL3), anti-CD11c (N418), anti-I-A/I-E (M5), anti-CD25 (PC61) all purchased from Biolegend. Splenocytes were incubated for 30mins at 4°C with the previously mentioned mAbs, washed with PBS then incubated with streptavidin magnetic beads for 20mins at 4°C. Negative fraction containing CD4 T cell was then stained with Viability Dye-APC-eF780, anti-TCRβ−APC (H57-597), anti-CD4-FITC (GK1.5), anti-CD8-PerCP-Cy5.5 (53-6.7) and anti-CD45RB-PE (C363-16A) mAbs. Live, TCRβ^+^, CD4^+^, CD45RB^high^ colitogenic T cells were then separated by fluorescent cell sorting using a BD FACS Aria II SORP with a purity over 90%. Colitis was induced in 9- to 11-week-old Rag1^−/−^ mice by retro-orbital injection of 1×10^6^ naive CD4^+^CD45RB^high^ from WT or Ets1-SE^−/−^ colitogenic CD4^+^ T cells in 100 μl of PBS. Weight loss of Ets1-SE^−/−^ or WT-injected Rag1^−/−^ was recorded every week for 6 to 7 weeks prior to mice euthanasia. Cell infiltration characterization, restimulation and macroscopic scoring was performed on week 6 to 7.

#### Macroscopic analysis

Macroscopic colonic tissue damage was evaluated by the Comparative Pathology Core (CPC) at the Veterinary school of the University of Pennsylvania using a scale described in Fig 3e. Colonic tissue specimens were excised 2 cm proximal to the cecum and immediately transferred into 10% formaldehyde to be embedded in paraffin. Colonic sections were then stained with hematoxylin and eosin by the Comparative Pathology Core (CPC) at the Veterinary school of the University of Pennsylvania. Each slide was scored (blind readings) by a single pathologist. Colon length was measured using a ruler right after mice euthanasia (Fig. S3a).

#### Colon Lamina propria (cLP) harvest digestion and cell infiltration phenotyping

Colons were excised and fat was removed using forceps, unrolled, and measured using a ruler. Colons were flushed with cold PBS to remove feces then were opened lengthwise and tissues were shaken in a petri dish containing cold PBS to remove any remaining feces. Colons were washed 2 times 10mins at 180 RPM and 37°C in PBS containing FBS (2%), HEPES (20mM) and EDTA (10mM). Colons were thoroughly washed 2 times with ice cold PBS and minced into 1 cm pieces using scissors. The minced colons were then digested PBS containing FBS (2%), HEPES (20mM), Collagenase D (1mg/ml), DNAse I (0.2mg/mL), and Dispase (0.1 U/ml). The digested colons were filtered through a 100μm cell strainer and pelleted. Cells were then enriched over an 80/40% Percoll gradient prior to staining or *ex vivo* restimulation. For cytokines production by colonic CD4^+^ T cells, cells were plated in a round bottom 96-well plate in complete RPMI medium containing PMA (Sigma; final concentration 100ng/mL) and Ionomycin (Sigma; final concentration 10ng/mL) and GolgiPlug (1X - BD Bioscience) and incubated for 4 hours at 37°C 5% CO_2_. Cells were then harvested and used for subsequent flow cytometry staining and analysis.

### Genomics and sequencing experiments

#### RNA-seq

Around 100,000 cells were washed once with 1x PBS before resuspending pellet in 350 μL Buffer RLT Plus (QIAGEN) with 1% 2-Mercaptoethanol (Sigma), vortexed briefly, and stored at −80°C. Subsequently, total RNA was isolated using the RNeasy Plus Micro Kit (QIAGEN). RNA integrity numbers were determined using a TapeStation 2200 (Agilent), and all samples used for RNA-seq library preparation had RIN numbers greater than 9. Libraries were prepared using the SMARTer® Stranded Total RNA-seq Kit v2-Pico Input Mammalian kit (Takara). Two technical replicates were generated for each experiment. Libraries were validated for quality and size distribution using a TapeStation 2200 (Agilent). Libraries were paired-end sequenced (38bp+38bp) on a NextSeq 550 (Illumina) or 61bp+61bp on Novaseq 6000 (Illumina).

#### ATAC-seq

ATAC-seq was performed as previously described with minor modifications^10^. Fifty thousand cells were pelleted at 550 g and washed with 50 μL ice-cold 1x PBS, followed by treatment with 50 μL lysis buffer (10 mM Tris-HCl [pH 7.4], 10 mM NaCl, 3 mM MgCl2, 0.1% IGEPAL CA-630). Nuclei pellets were resuspended in 50 μL transposition reaction containing 2.5 μL Tn5 transposase (FC-121-1030; Illumina). The reaction was incubated in a 37°C heat block for 45 min. Tagmented DNA was purified using a MinElute Reaction Cleanup Kit (QIAGEN) and amplified with varying cycles, depending on the side reaction results. Libraries were purified using a QIAQuick PCR Purification Kit (QIAGEN). Libraries were validated for quality and size distribution using a TapeStation 2200 (Agilent). Libraries were paired-end sequenced (38bp+38bp) on a NextSeq 550 (Illumina) or 61bp+61bp on Novaseq 6000 (Illumina).

#### Hi-C

Hi-C libraries were generated on 1×10^6^ cells using with Arima-HiC+ kit (Arima Genomics) and Accel-NGS @S Plus DNA Library kit (21024 Swift Biosciences), according to the manufacturer’s recommendations. Libraries were validated for quality and size distribution using Qubit dsDNA HS Assay Kit (Invitrogen, cat# Q32851) and TapeStation 2200 (Agilent). Libraries were paired-end sequenced (66bp+66bp) on NovaSeq 6000 (Illumina).

#### CUT&RUN

CUT&RUN was performed on in-vitro polarized Th1 cells using CUTANA™ ChIC/CUT&RUN Kit (EpiCypher, Cat#14-1048), using manufacturer’s recommendation. Briefly, 4 × 10^5^ live cells were sorted out and nuclei were extracted, washed, and allowed to adsorb onto activated ConA beads. Cells were then resuspended in recommended buffer, 0.5 mg of antibody was added, mixed well, and allowed to incubate at 4°C overnight on a nutator. Recommended antibodies were used, including anti-H3K27ac (Acetyl-Histone H3 (Lys27) (D5E4) XP® Rabbit mAb, CST, Cat #8173S), anti-ETS-1 (C-20, SantaCruz, Cat# sc-350X) and anti-CTCF (Millipore, Cat# 07-729), along with positive and negative controls. Subsequently, the reactions were washed with cell permeabilization buffer and incubated with pAG-MNase, and the DNA was isolated for the antibody-bound regions. At least two biological replicates were generated for each experiment. Library preparation was carried out using NEBNext Ultra II DNA Library Prep Kit for Illumina (NEB) and were paired-end sequenced (38bp+38bp) on a NextSeq 550 (Illumina) or 61bp+61bp on Novaseq 6000 (Illumina).

### Oligopaint FISH probe generation

The OligoMiner pipeline was used to design Oligopaint libraries as performed earlier^10^. 42bp probes were designed to a 50 kbp region at a density of approximately 5 probes per kilobase for the *Ets1* locus using the GRCm38.87 genome.

### Oligopaint FISH hybridization

Primary naïve CD4^+^ T cells isolated from spleen were activated overnight on plates coated with 2 μg/mL of anti-CD3, using complete RPMI media^10^, supplemented with 1 ng/mL of recombinant IL-2 and 2 ug/mL of soluble anti-CD28. Following this, the cells were diluted to 4 million cells per mL, and 80uL of diluted cells were added to polysine microscope slides (Thermo Scientific, cat#P4981-001) using silicone isolators (Electron Microscopy Sciences, cat #70339-05). Cells adhered to the slides for 1 hour at room temperature inside humidified chambers. Cells were then briefly washed in 1xPBS, fixed in 4% formaldehyde (Fisher Scientific, cat#PI28908) in PBS for 10 min, and then washed in 1xPBS. Slides were stored temporarily in 1xPBS at 4°C or used immediately for DNA FISH. Cells were permeabilized in 0.5% Triton in PBS for 15 min and dehydrated with an ethanol row of 70%, 90%, and 100% ethanol for 2 min each. After allowing the slides to dry for 3-5 minutes, cells were washed in 2XSSCT/50% formamide (0.3M NaCl, 0.03M sodium citrate, 0.1% Tween-20) at room temperature for 5 minutes, 2.5 min at 92°C in 2XSSCT/50% formamide, and 20 min at 60°C in 2XSSCT/50% formamide. For primary probe hybridization, slides were cooled down to room temperature, and cells were immersed in hybridization buffer (10% dextran sulfate, 50% formamide, 4% PVSA, 5.6 mM dNTPs, and 10ug of RNase A) containing 50 pmol of primary Oligopaint probes, covered with a coverslip (Fisher Scientific, cat#12-548-5M), and sealed with no-wrinkle rubber cement (Elmer’s). Cells were denatured for 2.5 min at 92°C on top of a heated block, followed by hybridization at 37°C in a humidified chamber for ∼16 hrs. Coverslips were then carefully removed using a razor blade, and cells were washed for 15 min in 2XSSCT at 60°C, followed by 10 min wash at room temperature in 2XSSCT shaking at 75 rpm and another 10 min wash at room temperature in 0.2XSSCT. After allowing the slides to air-dry, cells were immersed in secondary hybridization buffer (10% dextran sulfate, 10% formamide, and 4% PVSA) with 2pmol bridges and 10pmol of secondary probes (Alexa-488, Atto-565, and Alexa-647), covered with a coverslip (Fisher Scientific, cat#12-548-5M), and sealed with no-wrinkle rubber cement (Elmer’s). Slides were then incubated in the dark in a humidified chamber for 2 hrs at room temperature. Coverslips were then carefully removed using a razor blade, and slides were briefly washed in 2XSSCT at room temperature, followed by a 5 min wash in 2XSSCT at 60°C, a 5 min wash in 2XSSCT with DAPI (0.1 µg/mL), and a 5 min wash in 0.2XSSC. Slide were held in 2XSSC before mounting with Slowfade Gold Antifade Reagent (Invitrogen by Thermo Fisher Scientific, cat#S36936) and sealing with Sally Hansen’s “dries instantly top coat”.

### Genomics Data Analysis

#### HiChIP data processing and 3D clique analysis

H3K27ac HiChIP measurements in mouse DP T cells were generated in our previous study^10^. Data analysis was performed as previously described: Raw reads for each HiChIP sample were processed with HiC-Pro (version v2.5.0)^63^ to obtain putative interactions with default parameters except LIGATION_SITE = GATCGATC and GENOME_FRAGMENT generated for MboI restriction enzyme. Valid pairs (VI), self-circle (SC) and dangling-end (DE) interactions in cis were used as input for significant interaction calling in ‘.bedpe’ format. Mango (version 1.2.0) (Phanstiel et al., 2015) step 4 identified putative significant interaction anchors by MACS peak calling with MACS_qvalue = 0.05 and MACS_shiftsize = 75. Mango step 5 identified significant interactions with default parameters except maxinteractingdist = 2000000 and MHT = found. Two biological repeats for each strain were processed and only significant interactions with PETs >= 2 reproduced in both replicates were used for further analysis. For each library, each significant interaction was normalized to contacts per hundred million, i.e., divided by the number of interactions in the Mango input.bedpe file and multiplied by 1E8. Mango outputs of two biological replicates where two anchors were within 5kbp were called reproducible interactions in DP T cells.

#### 3D clique analysis

3D clique analysis was performed following the same procedure as reported earlier^9,10^. An undirect graph of regulatory interactions was constructed for reproducible interactions with at least one H3K27ac peak at one anchor. In this graph, each vertex was an enhancer or a promoter and each edge was a significant and reproducible enhancer-enhancer, enhancer-promoter, or promoter-promoter interaction. “3D Cliques” were defined by spectral clustering of the regulatory graph interactions using cluster_louvain function in igraph R package with default parameters. A 3D clique connectivity was defined as the number of edges connecting vertices within the clique. The connectivity of cliques was ranked in ascending order and plotted against the rank. The cutoff for hyperconnected 3D cliques was set to the elbow of the curve and a tangent line at the cutoff was shown. Super-enhancers were defined using H3K27ac ChIP-seq in DP T cells as described earlier^36^. Annotation of noncoding RNA was performed using gencode.vM10.annotation.gtf file. Architectural stripes were defined using Stripen^14^n using Hi-C measurements in DP T cells. Odds ratio analysis was performed to evaluate the significance of enrichment of super-enhancers, ncRNA and architectural stripes.

### Hi-C data analysis

#### Hi-C alignment

We used the ArimaHiC protocol to generate our Hi-C libraries. Hence, we followed the manufacturer’s recommendations and processed the data with HiC-Pro using parameters “LIGATION_SITE=GAATAATC,GAATACTC,GAATAGTC,GAATATTC,GAATGATC,GACTAATC,GACTACTC,GA CTAGTC,GACTATTC,GACTGATC,GAGTAATC,GAGTACTC,GAGTAGTC,GAGTATTC,GAGT GATC,GATCAATC,GATCACTC,GATCAGTC,GATCATTC,GATCGATC,GATTAATC,GATTACT C,GATTAGTC,GATTATTC,GATTGATC” and GENOME_FRAGMENT file was generated using “digest_genome.py -r ^GATC G^AATC G^ATTC G^ACTC G^AGTC”. ValidPairs generated by HiC-Pro were further converted to cool and hic files.

#### Compartment analysis

The principal component analysis (PCA) was performed with 50 kbp resolution for both wild type and *Ets1*-SE^−/−^ Th1 Hi-C data. To generate the PC1 plot in the Figure S6a, a customized script (cool_compartment.py) utilizing *cooltools* was used.

#### TAD analysis

TAD coordinates were estimated using two Perl scripts named ‘*matrix2insulation.pl*’ and ‘*insulation2tads.pl*’ from the cworld-Dekker Github page for both wild type and *Ets1*-SE^−/−^ Th1 Hi-C data. The overlap of TAD boundaries was evaluated using the *findOverlaps* function in an R package called GenomicRanges.

#### Loop analysis

Loops were called using Mustache^60^ from both wild type and *Ets1*-SE^−/−^Th1 and Th2 Hi-C with 5kbp resolutions. Loops with FDR < 0.1 were used for further analysis. The scatter plot for loop intensity of wild type and *Ets1*-SE^−/−^ mice was generated, and the loops with intensity higher than |WT/ *Ets1*-SE^−/−^ |>0.5 was highlighted.

#### Triangle heatmaps

Triangle heatmaps for 3D chromatin conformation data and corresponding tracks were generated using Sushi R package (version 1.28.0)

#### CUT&RUN data analysis

The FASTQ files of CUT&RUN experiments were aligned to the bam file using BWA (version 0.7.17-r1188). In this process, minor chromosomes such as mitochondrial chromosome or chrY were removed using samtools (version 1.11). Next, duplicated reads were removed using Picard (version 2.26.7) and then the bam files were indexed using samtools. BigWig files were generated using bamCoverage (version 3.3.2) with parameters ‘normalizedUsing=CPM, binsize=30, smoothLength=300, p=5, extendReads=200’. For peak calling, macs2 (version 2.1.4) was used with following commands: ‘macs2 callpeak -t input_file - c control -g mm -n output_path –nomodel -f BAMPE -B –keep-dup all –broad –broad-cutoff 0.25 -q 0.25’. For the background (control), the bam file of IgG CUT&RUN data was used. CUT&RUN peaks from two conditions and both replicates were merged and the number of fragments in each peak were counted with bedtools. The count data of each peak was then fed to DESeq2 for differential analysis.

### Integration of CUT&RUN data with 3D loop analysis

#### Loop classification

The loops called from the wild type Th1 Hi-C with 5kb resolution were classified based on the occupancy of CTCF and/or ETS1. First, the CTCF and ETS1 peaks from replicates were combined, respectively. Next, both ends of each loop (loop anchors) were extended by ±5 kbp and then the occupancy of CTCF or ETS1 peaks within loop anchors was examined using the R package named GenomicRanges.

#### DESeq2 analysis and pileup plot

To identify CTCF and ETS1 peaks that were lost or gained in a coordinated manner in Ets1-SE^−/−^ Th1 cells, DESeq2 was performed by (1) combining CTCF and ETS1 peaks across all replicates, (2) counting reads across this union of peaks, and (3) comparing wildtype and Ets1-SE^−/−^ Th1 cells samples regardless of the protein occupancy. As a result, 988 and 364 co-bound CTCF-ETS1 peaks were detected to be co-lost and co-gained, respectively (|log2 fold change| > 1 and p-value<0.05). 1D heatmaps using deeptools or pileup plots for 3D interactions using coolpuppy examined the 1D and 3D features of these peak sets. For coolpup.py, a parameter –pad=50 was used. Local pileup analysis at different sets of peaks was done with coolpup.py 63 using parameters “--pad 250 --local”. The average interaction of the upright corner of pileup plots is quantified with custom python script, by parsing the results from coolpup.py. Interactions between regions in bedfiles were done with parameters “--mindist 200000 --maxdist 2000000”. Pileup of loops was also plot with coolpup.py. To do multiple pileup analysis parallelly, we used GNU parallel to run the shell script.

#### Deeptools analysis of CUT&RUN and ATAC-seq data

The differentially gained or lost sites in Naive CD4+ T, Th1 and Th2 cells were obtained using DESeq2 (|log2 fold change| > 1 and adjusted p-value<0.05) and then combined using bedtools sort and merge function. Next, deeptools plot was generated with computeMatrix function using following parameters: reference-point –referencePoint center -a 2000 -b 2000. The heatmap was generated with the ‘plotHeatmap’ function with the following parameters: ‘–heatmapHeight 15 –averageTypeSummaryPlot median –colorMap Greys’. In comparison of Naive CD4 T and Th1 cells, we performed DESeq2 for Naive CD4 T and Th1 cells (Ets1-SE+/+). Then, Naive CD4 T-specific (log2 fold change > 1 and adjusted P-value < 0.05), Th1-specific (log2 fold change < -1 and adjusted P-value < 0.05) and non-significant (adjusted P-value > 0.05) peaks were obtained and fed to computeMatrix/plotHeatmap function.

#### RNA-seq data analysis

The FASTQ files of RNA-seq experiments were aligned and further counted using STAR 2.7.7a with parameters ‘--outSAMtype BAM SortedByCoordinate -- outWigType wiggle read1_5p --outWigStrand Stranded --outWigNorm RPM --quantMode GeneCounts’. Next, DESeq2 was performed to identify differentially expressed genes (|log2 fold change|>1 and adjusted p-value < 0.05).

#### ATAC-seq data analysis

The alignment process of ATAC-seq data is identical to that of CUT&RUN data except MACS2 parameters which is as follows: ‘macs2 callpeak -t input_file -g mm -n output_path –nomodel -f BAM -B –keep-dup all –broad –broad-cutoff 0.1 -q 0.1’. ATAC-seq peaks from two conditions and both replicates were merged and the number of fragments in each peak were counted with bedtools. The count data of each peak was then fed to DESeq2 for differential analysis.

#### GSEA and Gene Ontology analysis

Pre-ranked Gene Set Enrichment Analysis (GSEA) was used for enrichment analyses. Gene-sets were provided by DESeq2 for downregulated and upregulated genes in *Ets1*-SE^−/−^ compared with wildtype Th1 cells. Pre-ranked genes based on DESeq analysis of across different conditions were used. Metascape using ImmuneSigDB was used for gene-ontology analysis.

#### ImmGen gene expression data processing

Gene Skyline feature for RNA-seq data provided by ImmGen was used to profile Ets1 and Fli1 expression levels across various cell types.

#### Motif analysis

To find the conserved motifs from the lost or gained ATAC-seq/Hi-C peaks(=loops) in *Ets1*-SE^−/−^ mice, a function named ‘findMotifsGenome.pl’ in Homer program (version 4.11) was utilized. For ATAC-seq data of naive CD4 T cells, gained and lost peaks in *Ets1*-SE^−/−^ mice were determined by DESeq2 results such that |log2 fold change| > 1 and adjusted P-value < 0.05. Here, the peaks that were not significantly changed (adjusted P-value>0.8) were used as background. The loops gained and lost in Ets1-SE were searched from Hi-C data of Th1 cells. Here, gained and lost loops were defined as those |fold change|>0.5, and the loops showing minute changes (|fold change| < 0.1) were considered as background. These loop coordinates were then fed to ‘bedtools intersect (version 2.30.0)’ to obtain the coordinates overlapping with CUT&RUN peaks of ETS1.The output consists of (1) consensus motifs (known motifs) and (2) de novo motifs conserved in the input sequences (homer motifs). Here, we reported the latter one.

#### Statistical analysis

For all experiments, the difference between two groups was calculated using unpaired t-test (Mann and Whitney) Prism 10 GraphPad Software. ANOVA and Bonferroni test were used for multiple comparisons (ns = not significant, * = p ≤ 0.05, ** = p ≤ 0.01, *** = p ≤ 0.0005, **** = p ≤ 0.0001). The Mann and Whitney unpaired t-test was used for comparisons of two conditions within one group. One-way ANOVA was used to compare more than two conditions for one group. The Two-way ANOVA was used to compare two conditions across multiple groups. All graphs show the mean and the standard error of the mean (SEM). One- and Two-way ANOVA were corrected for multiple comparison using Bonferroni correction.

### Oligopaint FISH imaging and analysis

Imaging was carried out on a Bruker Vutara VXL in the Widefield imaging modality, which has an imaging field-of-view (FOV) of 200 µm x 200 µm. The VXL has a 60X silicon oil immersion objective with a 1.3 numerical aperture (NA). Z-stack size was maximum 20 µm with a Z-step size of 150 nm. Analysis was carried out on the raw images in a semi-automated manner on a cell-by-cell basis as describe in Raj. et al. 2008 (https://github.com/arjunrajlaboratory/rajlabimagetools). Briefly, the DAPI signal was used for manual nuclei segmentation of activated naïve CD4 T cells. The exact numbers of nuclei analyzed per genotype are as follows: 561 nuclei for wildtype and 727 nuclei for the cells from *Ets1-SE*^−/−^. One mouse per genotype was used. Spots for each of the 3 channels (Alexa-488, Atto-565, and Alexa-647) were individually detected using a linear filter approximately conforming to a Laplacian convolved with a Gaussian. For each spot, the brightest z slice was used as the z coordinate. Centroid positions for each spot in xy were found by fitting a Gaussian. X, Y, and Z coordinates were extracted, and pairwise Euclidean distances between nearest neighbors were calculated. The spatial distance of two probes in each cell were determined by taking a minimum from all pair-wise distances of corresponding probes. The Kolmogorov-Smirnov test was used to compare differences in the cumulative distributions, and the Mann-Whitney test was used to compare differences in the medians. In addition, the proportion of cells having probe distance less than threshold was examined for 3D clique dynamics analysis.

### Representative Image processing

Imaging was carried out on a Leica Multiphoton Confocal using a 63X oil immersion objective with a 2.0 zoom factor, a pixel size of 58.77 nm x 58.77 nm, and Z-stack size of 15 µm with a Z-step size of 300 nm. Z-stacks were maximally projected. Each cell, allele, and locus for each strain were individually processed using ImageJ via adjusting the brightness/contrast/minimum/maximum, as well as smoothing.

## Acknowledgments

We thank helpful discussions with members of the Vahedi lab in addition to Robert B. Faryabi, Yeqiao Zhou, Kenneth Zaret, Rajain Jain, Sara Cherry, Arjun Raj, and E. John Wherry. We would like to thank Behdad Afzali and his lab for help and discussion about the CUT&RUN experiments in T helper cells. The work in this manuscript was supported by the Burroughs Welcome Fund, the Chang Zuckerberg Initiative Award, and the NIH grant R01AI168240 (J. H-M. and G.V.); W. W. Smith Charitable Trust, the Penn Epigenetics Institute, the Sloan Foundation, and the NIH grants UC4DK112217, U01DK112217, R01HL145754, U01DK127768, U01DA052715 (G.V); the PEW Charitable Trust and the NIH grant R01HL136572 (J. H-M.); and the NIH grant R01 AI106352 (B. L. K.).

